# Automethylation of lysine methyltransferase SETDB1 on H3K9-like motifs regulates interactions with chromodomain proteins and controls its functions

**DOI:** 10.1101/2025.10.22.683908

**Authors:** Paola Cruz-Tapias, Guillaume Velasco, Roberta Rapone, Laurence Del Maestro, Victor Cochard, Guillaume Chevreux, Véronique Joliot, Ekaterina Boyarchuk, Slimane Ait-Si-Ali

## Abstract

Histone H3 lysine 9 trimethylation is essential for heterochromatin formation and maintenance, genome stability, and silencing of transposable elements (TEs) in embryonic stem cells (ESCs). The H3K9-specific lysine methyltransferase SETDB1 is crucial for mammalian development, as it controls ESC pluripotency and viability by regulating gene expression and TE silencing. Here, we demonstrate that SETDB1 undergoes automethylation on two lysine residues located within H3K9-like motifs in its catalytic domain. Notably, these automethylated lysines are necessary for the normal growth and viability of ESCs. While SETDB1 automethylation does not affect its catalytic activity, it is crucial for its interaction with chromodomain-containing partners, including SUV39H1, CDYL, and Heterochromatin Protein 1 gamma (HP1gamma). The integrity of the two automethylated lysines is required for SETDB1 localization to its target sites and for effective silencing of both coding genes and TEs. Expression of an automethylation-deficient SETDB1 fails to properly establish H3K9me3 and disrupts HP1gamma recruitment at target loci. Collectively, our findings uncover a previously unknown mechanism regulating SETDB1 function, essential for maintaining the fitness of mouse ESCs.

**Highlights:**

– SETDB1 undergoes automethylation on two lysines within its catalytic domain
– The automethylation-deficient form of SETDB1 is enzymatically active, yet it compromises mESCs fitness
– SETDB1 automethylation-deficient mutant impairs chromatin association and pan-genomic H3K9me3 landscape
– SETDB1 automethylation regulates its interactions with many chromodomain-containing partners

## INTRODUCTION

Lysine methylation (Kme) is a stable post-translational modification (PTM) that regulates protein-protein interactions, but also protein stability, activity, or subcellular localization. In the nucleus, Kme also regulates gene expression by modifying both histone and non-histone proteins (Hamamoto et al., 2015). Kme is added by lysine methyltransferases (KMTs), removed by demethylases (KDMs), and can recruit effector proteins via reader domains like chromodomains and Tudor domains, impacting diverse cellular processes.

Heterochromatin is a conserved structural feature of eukaryotic genomes that plays a central role in genome organization, gene expression regulation, and genomic stability (Allshire and Madhani, 2018; Grewal, 2023). It is characterized by high levels of histone H3 lysine 9 di- and tri-methylation (H3K9me2/3). Heterochromatin formation initiates at defined nucleation sites through the targeted recruitment of silencing complexes, followed by lateral spreading to establish a repressive chromatin domain (e.g., H3K9me2 or H3K9me3 domains). This spreading is driven by H3K9-specific methyltransferases, which recognize pre-existing methylation marks via their methyl-lysine-binding domains and thereby reinforce H3K9 methylation in a positive feedback loop. Additional reader proteins, such as Heterochromatin Protein 1 (HP1) and CDYL, further stabilize and extend these silenced domains by binding H3K9me2/3 and promoting chromatin compaction and heterochromatin maintenance (Allshire and Madhani, 2018; Grewal, 2023).

H3K9me3 is essential for heterochromatin formation and silencing of transposable elements (TEs) in mouse ESCs (mESCs) (Padeken et al., 2022). This histone mark is established through sequential, processive addition of methyl groups by the SUV39 KMT family, which includes G9A/GLP, SETDB1, and SUV39H (comprising two paralogues, H1 and H2). All family members contain a conserved catalytic domain named SET along with other functional domains. Notably, methyl-lysine-binding modules, such as the ankyrin repeats on G9A/GLP, the Tudor domains on SETDB1, and the chromodomain on SUV39H (Mozzetta et al., 2015), enable these enzymes to act as both writers and readers of lysine methylation.

G9A and GLP are responsible for H3K9 mono- and dimethylation mainly in euchromatin, establishing large domains of H3K9me2 in ESCs and providing methylated substrates for other H3K9 KMTs (Mozzetta et al., 2015; Padeken et al., 2022). Deletion of G9A and GLP genes is lethal during early embryonic development at 9.5 days post-coitum (dpc), reflecting their essential role in regulation of gene expression (Tachibana et al., 2002), but also in repression of certain TE families, such as murine endogenous retrovirus-L (MERVL) and Intracisternal A-particle (IAP) sequences (Maksakova et al., 2013).

SUV39H1 and SUV39H2 catalyze H3K9 di- and tri-methylation mainly at pericentromeric repeats and telomeric constitutive heterochromatin (Lehnertz et al., 2003), but are also involved in H3K9me3 establishment at several endogenous retroviral elements (ERVs) and long interspersed nuclear element (LINE) LINE-1 (Bulut-Karslioglu et al., 2014). SUV39H trimethylates an already mono- or di-methylated H3K9 by G9A/GLP or SETDB1 (Garcia-Cao et al., 2004; Gauchier et al., 2019). Unlike *G9A/GLP* or *SETDB1* knock-out (KO), double *SUV39H1* and *SUV39H2* KO mice showed no developmental delay or defect but a high post-natal lethality mainly due to genome instability (Peters et al., 2001).

SETDB1 is crucial for mammalian development, as it maintains ESC pluripotency in the early embryo by silencing numerous TE classes. *SETDB1* gene KO in mice is lethal during early development at the peri-implantation stage at 3.5-5.5 dpc (Dodge et al., 2004). SETDB1 is essential for the survival of ESCs and it influences ESCs identity, pluripotency and self-renewal (Bilodeau et al., 2009; Yeap et al., 2009). SETDB1 is capable of establishing all three H3K9 methylation levels and contains Tudor domains that facilitate binding to methylated lysines and enable protein-protein interactions with co-factors (Chandrasekaran et al., 2024), such as with the Activating Transcription Factor 7-interacting protein (ATF7IP), HP1, and KAP1 (Schultz et al., 2002; Timms et al., 2016). These interactions regulate SETDB1’s targeting and activity. Specifically, ATF7IP influences SETDB1 enzymatic activity, stability, and subcellular localization, directs it to TEs in ESCs, and enhances its activity towards H3K9 trimethylation (Tsusaka et al., 2019). Conversely, the SETDB1/HP1 interaction directs SETDB1 towards pericentromeric chromatin to establish H3K9 monomethylation (Loyola et al., 2009; Tsusaka et al., 2019). Beyond targeting, HP1 binding also promotes spreading of H3K9 methylation (Bannister et al., 2001; Lachner et al., 2001).

H3K9 KMTs can cooperate at specific sites. SETDB1 forms a functional complex with other H3K9 KMTs, such as G9A, GLP, and SUV39H1, to facilitate transcriptional silencing (Fritsch et al., 2010). In the mouse ESCs, both G9A and SETDB1 are crucial for the complete silencing of ERVs (Maksakova et al., 2013). SETDB1 and SUV39H also cooperate and share functions; they target common elements, including TEs like LINEs and ERVs (Bulut-Karslioglu et al., 2014), and coordinate the propagation of H3K9me3 in pericentromeric heterochromatin, telomeric repeats (Gauchier et al., 2019), and at nuclear peripheral heterochromatin (Loyola et al., 2009; Towbin et al., 2012).

Additionally, beyond H3K9, these KMTs can methylate non-histone substrates and undergo automethylation. For example, SETDB1 methylates several non-histone substrates, including ING2, AKT, and p53 proteins (Binda et al., 2010; Fei et al., 2015; Guo et al., 2019). Remarkably, some non-histone proteins methylated by H3K9 KMTs carry histone mimic motifs that mediate protein–protein interactions (Chin et al., 2007; Sampath et al., 2007; Tsusaka et al., 2018). Thus, G9A can methylate its own histone mimic motif to promote HP1 binding (Chin et al., 2007; Sampath et al., 2007; Tsusaka et al., 2018). G9A/GLP also methylate SETDB1 cofactor ATF7IP at a histone mimic motif that mediates its interaction with the chromodomain-containing protein MPP8 (Tsusaka et al., 2018). Finally, SUV39H ortholog in *Schizosaccharomyces pombe*, Clr4, is automethylated on several lysine residues (Iglesias et al., 2018; Sampath et al., 2007), modifications that regulate its activity and/or interaction with protein partners.

Here, we identify a novel regulatory mechanism whereby SETDB1 undergoes automethylation on two conserved lysines within its catalytic domain, embedded in H3K9-like histone-mimic motifs. The integrity of these methylated lysines is essential for the fitness and viability of mESCs. Notably, SETDB1 automethylation does not influence its catalytic activity, but modulates its interactions with chromodomain-containing partners, including HP1γ, CDYL, and SUV39H. Maintaining the integrity of the two methylated lysines in SETDB1 is crucial for its enrichment on chromatin and efficient H3K9me3-mediated silencing of both coding genes and TEs. Collectively, our findings uncover a previously unknown mechanism regulating SETDB1 functions in mESCs.

## RESULTS

### SETDB1 undergoes automethylation *in vivo*

A lysine methylome analysis performed by mass spectrometry (MS) in human cells showed that SETDB1 is extensively methylated, including within its own catalytic domain on lysines 1170 and 1178 (Guo et al., 2014). Since the two methylated lysines are embedded within an H3K9-like ARKS motif (**Figure 1A**), which is the preferential SETDB1 substrate, we checked if this methylation results from automethylation activity. Thus, to assess whether SETDB1 is capable of methylating itself as a non-histone substrate, we performed *in vitro* methylation assays using recombinant proteins. We first produced different human SETDB1 truncated mutants as GST-fusion proteins (**Figure 1B**). We used recombinant active full-length (FL) human SETDB1 enzyme and radioactive S-adenosyl-L-[methyl-^3^H]-methionine (^3^H-SAM) as methyl donor group. Strikingly, the upper band detected in the reactions with the truncated SETDB1 mutants revealed automethylation on full-length SETDB1 (**Figure 1C**). Interestingly, when histone H3 was provided as substrate, SETDB1 displayed a clear preference for methylating activity H3 over itself (**Figure 1C**). Notably, the bifurcated domain and the carboxy-terminal region (C-ter) of SETDB1 were methylated by the FL SETDB1, in contrast to the pre-SET and post-SET domains (**Figure 1C**). This indicates that SETDB1 automethylation events take place within the C-terminal part of the bifurcated domain.

**Figure 1.**
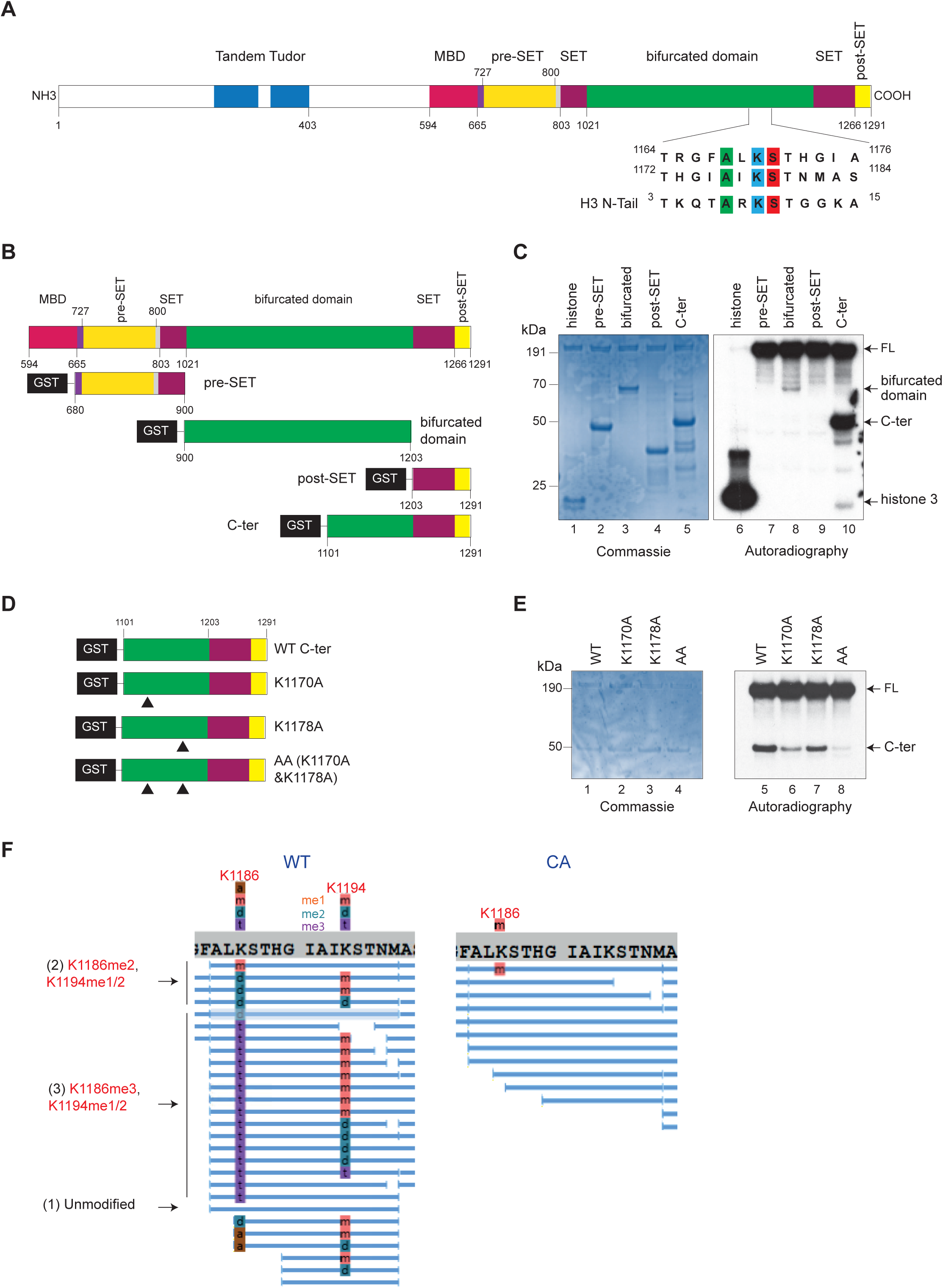
SETDB1 undergoes auto-methylation on lysines K1170 and K1178 *in vitro* and is methylated in the cells. **A.** Schematic representation of human SETDB1 protein. Tudor: Tudor domain; MBD: methylated DNA binding domain; SET: SET domain. Numbers correspond to amino acid position. Lower: sequence alignment of the region surrounding the SETDB1 K1170 and K1178 methylation sites and the sequence surrounding H3K9 on histone H3 N-terminal tail (N-Tail). Sequence conservation across species is shown in Figure S1A. **B.** Schematic illustration of recombinant GST-fused SETDB1 deletion mutants used in (C). **C.** Radioactive methylation assay performed on recombinant GST-SETDB1 fusion mutants, illustrated in (B), or GST alone as a negative control (not shown here), incubated with recombinant active full-length SETDB1. Purified GST-SETDB1 proteins were incubated with recombinant active SETDB1 in the presence of radioactive S-Adenosyl Methionine (SAM) and separated by SDS-PAGE. The gel was Coomassie-stained (left, lanes 1-5) and methylation was revealed by autoradiography (right, lanes 6-10). Full-length SETDB1 auto-methylation is observed in the upper part of the gel (lanes 7 to 10), in the absence of H3. As a positive control of SETDB1 activity, methylation of histone H3 within purified nucleosomes (lane 6) is shown. **D.** Schematic illustration of recombinant GST-SETDB1 deletion mutant encompassing amino acids 1101 to 1290 mutants used in (E), either wild-type (WT) or including the indicated single, or combined, lysine-to-alanine substitutions at the indicated positions (K1170A and K1178A). **E.** Radioactive methylation assay performed as in (C) to show SETDB1 auto-methylation on K1170 and K1178. **F.** Schematic representation of the abundance of the unmodified and modified peptides detected by mass spectrometry. The two most abundant modified peptides include dimethylation at K1186, and mono- and dimethylation at K1194 for one, and trimethylation at K1186 and mono/di-methylation at K1194 for the other.

We next used the GST-SETDB1 fusion construct that encodes the very carboxy-terminal region (amino acids 1101-1290) to generate three mutants harboring lysine (K)-to-alanine (A) substitutions in one or two putative methylation sites (K1170A and/or K1178A) located within “H3K9-like” A(L/I)KS motifs (**Figure 1D**) and performed *in vitro* radioactive methylation assays. Our results showed that the methylation level was substantially reduced in the GST-SETDB1 AA double-lysine mutant (**Figure 1E**), suggesting that K1170 and K1178 are required for SETDB1 automethylation.

Interestingly, the two automethylated lysines on human SETDB1 are highly conserved in many species, including in the mouse, where they correspond to lysines 1186 (K1186) and 1194 (K1194), respectively (**Figure S1A**). Therefore, we first checked if these automethylation events occur in mouse cells before studying their roles in mESCs, where SETDB1 is essential (Bilodeau et al., 2009; Dodge et al., 2004; Yeap et al., 2009) (see below).

Thereby, to assess both SETDB1 methylation and automethylation in mouse cells, we used *Setdb1* KO immortalized mouse embryonic fibroblasts (MEFs), in which SETDB1 expression was rescued using transiently transfected Myc-tagged wild-type (WT) or catalytic-dead mutant (C1243A point mutation, depicted as CA on the figures) SETDB1 versions (**Figure S1B**). WT and CA SETDB1 were immunoprecipitated in duplicate, and analyzed by tandem MS (**Figure S1B, S1C,** and **S1D**). In the WT SETDB1, K1186 and K1194 were found to be heavily modified by multiple methylation events, whereas in the CA mutant (which still harbors K1186 and K1194), almost no methylation was detected (**Figure 1F**). The peptide [Alanine 1184 - Methionine 1198] obtained after chymotrypsin digestion carries the two modified sites and was essentially found with different combinations of mono-, di- and tri-methylation events (me1, me2 and me3, respectively) (**Figure 1F**) as depicted on the representative mass spectra of the unmodified peptide, and of the two most abundant modified peptides, which included dimethylation at both K1186 and K1194 for one and trimethylation at K1186 and monomethylation at K1194 for the other (**Figure S1D**). These two modified peptides were ten times more abundant than the unmodified one (**Figure 1F**).

Finally, we confirmed SETDB1 automethylation in MEFs through an orthogonal approach. *Setdb1* KO MEFs were transfected with either empty vector or vectors expressing Myc-tagged WT, double lysine mutant (K1186 and K1194, referred to as AA in the figures), or catalytically inactive SETDB1 (CA). Nuclear extracts were used for immunoprecipitation (IP) with Myc-trap technology, and the precipitates were then analyzed by western blot (WB) using anti-pan-tri-methyl-lysine (panKme) antibody. Interestingly, the double lysine mutant SETDB1 extract showed a reduction in the level of lysine methylation at the SETDB1 molecular weight. Notably, for the SETDB1 catalytically inactive CA mutant, the decrease was more dramatic (**Figure S1E**), suggesting the presence of other potential automethylation events. Indeed, a recent study identified automethylation of human SETDB1 on its N-terminus (K489/K490), located in another H3K9-mimic motif (https://doi.org/10.1101/2025.01.21.634156). We also detected these automethylations at the corresponding residues in our MS analysis of mouse protein (not shown). The presence of these and other potential automethylation sites could explain the residual signal detected by anti-panKme antibodies in K1186A/K1194A double mutant.

Altogether, our data reveal that SETDB1 is methylated on two highly conserved lysines embedded within H3K9-like motifs, and this methylation is dependent on its own catalytic activity.

### The automethylation-deficient SETDB1 mutant retains enzymatic activity, yet it compromises the viability and growth of mESCs

We next asked whether the automethylation-deficient (K1186A and K1194A double mutant) form of SETDB1 retains enzymatic activity. To this end, we immunoprecipitated different Myc-tagged forms of SETDB1 from *Setdb1* KO MEFs (as shown in **Figure S1E**) and performed an *in vitro* methylation assay using the IP products as the enzyme, a cocktail of histones as substrate, and radioactive SAM as methyl donor group. We found that both WT and automethylation-deficient forms of SETDB1 showed comparable catalytic activities toward histone H3 (**Figure 2A**). As expected, the catalytically inactive (C1243A, or CA) mutant displayed significantly reduced activity (**Figure 2A**), thereby validating our IP-based assay. These results indicate that the catalytic activity of automethylation-deficient SETDB1 is not impaired.

**Figure 2.**
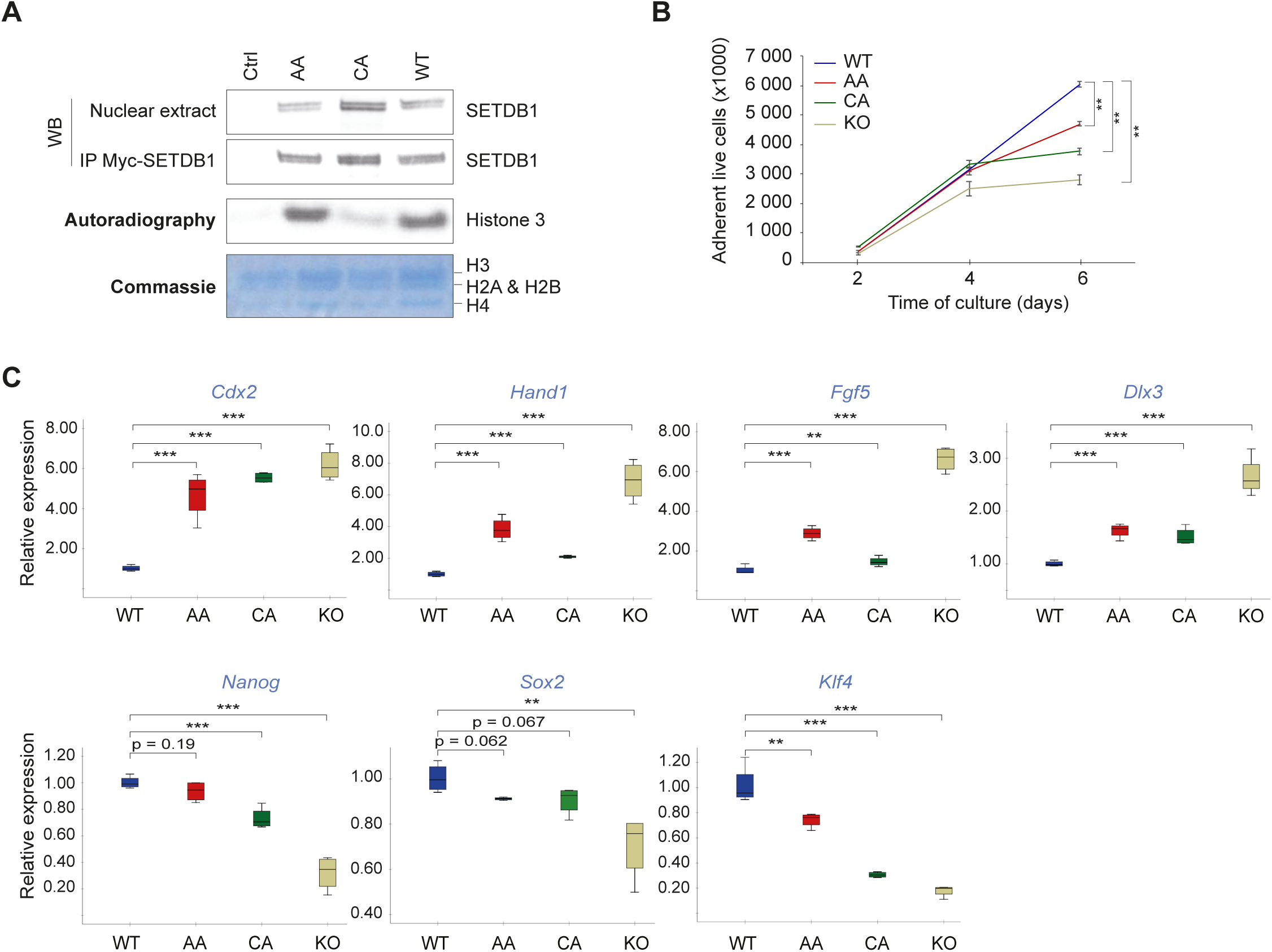
The automethylation-deficient SETDB1 mutant is enzymatically active, but it compromises mESCs’ fitness. **A.** Radioactive methylation assay performed on purified nucleosomes using immunoprecipitated SETDB1. Anti-Myc immunoprecipitation (IP) products from *Setdb1* KO iMEFs transfected with the empty vector (Ctrl) or expression vectors for the different SETDB1 versions (AA, CA or WT), were incubated with purified nucleosomes in the presence of radioactive SAM and separated by SDS-PAGE. Inputs (Nuclear extract) and IP products were analyzed by Western blot using Myc antibody. Methylation was revealed by autoradiography, and the nucleosomes by Coomassie staining. **B.** The automethylation-deficient SETDB1 mutant does not fully rescue growth defects in *Setdb1* KO ESCs. TT2 ESC line 33#06 harboring floxed *Setdb1* (Matsui et al., 2010) stably Flag-tagged wild-type (WT), automethylation-deficient (AA) or catalytically inactive (CA) SETDB1 were treated with tamoxifen (TAM) during 4 days, followed by 2 days without treatment. Endogenous SETDB1 was completely depleted by day four of TAM treatment (see Figure S2A). Adherent living cells were counted at the indicated times post-TAM treatment. Growth defect was observed at 6 days of culture for mESCs expressing the automethylation-deficient SETDB1 mutant. Cell viability was reduced at 4 days post TAM treatment for cKO *Setdb1* mESCs. For statistical significance, Student *t*-test was applied for data following normal distribution, otherwise, Mann–Whitney’s unpaired test was applied. (n = 3 biological replicates) **: p<0.05 **C.** Expression of automethylation-deficient (AA) or catalytically inactive (CA) SETDB1 mutants induce abnormal expression of differentiation genes (upper panel) and deregulates stemness genes such as *Klf4* (lower panel). ESC lines were treated as in (A), and gene expression was assessed by RT-qPCR using *Cyclophilin A* mRNA as the normalization control. Statistical significance was assessed using Student’s *t*-test for normally distributed data or the Mann–Whitney’s unpaired test otherwise (n = 4 biological replicates) **: p<0.05; ***: p <0.01.

SETDB1 is known to be essential for mESCs’ pluripotency, self-renewal, and survival (Bilodeau et al., 2009; Dodge et al., 2004; Yeap et al., 2009). Indeed, *in vitro*, *Setdb1* depletion in mESCs impairs cell proliferation and induces severe growth defects. We thus tested whether the presence of K1186/K1194 and the catalytic activity of mouse SETDB1 affect mESCs’ viability *in vitro*. To address this, we used conditional *Setdb1* KO TT2 mESCs (cKO) (Matsui et al., 2010), re-expressing either WT, catalytic-dead (CA) mutant, or the automethylation-deficient (double-lysine AA mutant) forms of SETDB1. In these TT2 mESCs, endogenous SETDB1 protein depletion starts at 48h (Rapone et al., 2023), and is achieved after 96h of treatment with 4-hydroxy-Tamoxifen (TAM) (**Figure S2A**). Importantly, SETDB1 protein levels were comparable in mESCs expressing WT and automethylation-deficient AA mutant SETDB1, while SETDB1 levels were higher in the cells expressing the catalytic-inactive CA mutant SETDB1 (**Figure S2A**). This could be attributed to the intrinsic activity of SETDB1, identified as a limiting factor in the expression of transgenes (Ngo and Puschnik, 2023), unlike the catalytically inactive form.

Next, we checked the distribution of SETDB1 in mESCs after TAM treatment, and, as shown in **Figure S2B**, SETDB1 is localized in the nucleus as well as in the cytoplasm in all cell lines expressing exogenous SETDB1, a phenotype comparable with the expression of the endogenous SETDB1 in non-treated parental mESCs (the 3306 TT2 cells, WT).

As previously reported by us and others (Matsui et al., 2010; Rapone et al., 2023), *Setdb1* cKO TT2 mESCs exhibit a growth defect after 6 days of TAM treatment. Notably, this growth defect was rescued by ectopic re-expression of WT SETDB1 in *Setdb1* cKO TT2 mESCs, whereas re-expression of the CA catalytically inactive SETDB1 mutant failed to rescue the defect (Matsui et al., 2010; Rapone et al., 2023). These observations prompted us to investigate whether the growth defect in *Setdb1* cKO TT2 mESCs could be rescued by re-expressing an automethylation-deficient AA form of SETDB1. To address this, we analyzed cell viability at 2, 4, and 6 days of TAM treatment. Our results revealed a decline in cell viability in *Setdb1* cKO mESCs, as expected, as well as in cells expressing the automethylation-deficient AA form, compared to those expressing WT SETDB1 starting at 4 days post-Tamoxifen treatment, although to a lesser extent than that observed with the catalytically-inactive CA mutant (**Figure 2B**). Moreover, this reduced proliferation was accompanied by a significant induction of apoptosis as measured by the Annexin V assay in cells expressing the automethylation-deficient AA SETDB1 mutant at 6 days, although the extent of apoptosis was lower than in KO cells (**Figure S2C**).

Finally, compared to WT mESCs, those expressing the AA automethylation-deficient SETDB1 express less stemness marker *Klf4* and more differentiation genes (*Cdx2*, *Hand1*, *Fgf5,* and *Dlx3*), though to a lower extent than in KO cells (**Figure 2C**). This indicates that the two methylated lysines are important for mESCs’ stemness phenotype.

Taken together, these results suggest that the expression of the automethylation-deficient form of SETDB1 compromises the fate and viability of mESCs after depletion of endogenous SETDB1, recapitulating, though less strongly, previously reported observations for the catalytically inactive form of SETDB1.

### Automethylation-deficient double-lysine SETDB1 mutant impairs chromatin association and H3K9me3 landscape, partially mimicking catalytic inactivation

We next investigated whether the integrity of K1186/K1194 in SETDB1 affects SETDB1 genomic occupancy and SETDB1-dependent H3K9me3 establishment. To address this, we performed genome-wide analyses for enrichment of Flag-tagged SETDB1 and H3K9me3 by ChIP coupled to high-throughput DNA sequencing (ChIP-seq) in mESCs expressing either the WT, the CA catalytically inactive, or the AA automethylation-deficient forms of SETDB1.

We were able to detect around ∼4000 SETDB1-binding sites in the cells expressing WT SETDB1 and ∼6900 for the SETDB1 catalytically inactive mutant (q<0.05), relative to 2754 genes and 3823 genes, respectively, while only 674 SETDB1-binding sites (q<0.05), relative to 550 genes were detected in the cells expressing the automethylation-deficient form of SETDB1 (**Figure 3A** left, and **3B**). Notably, SETDB1 protein levels were comparable in mESCs expressing WT SETDB1 and automethylation-deficient AA mutant SETDB1, while SETDB1 protein levels were higher in the cells expressing the CA catalytic-inactive mutant SETDB1 (**Figure S2A**). On the other hand, we detected 14500 H3K9me3-enriched regions, associated with 6460 genes in the cells expressing WT SETDB1, which is more than twice compared to cells expressing the automethylation-deficient AA SETDB1 mutant (∼5100 H3K9me3-enriched regions, corresponding to 3280 genes). Only 132 H3K9me3-enriched regions, relative to 100 genes, were detected in the cells expressing the SETDB1 CA catalytic inactive mutant (**Figure 3A** left, **3B**).

**Figure 3.**
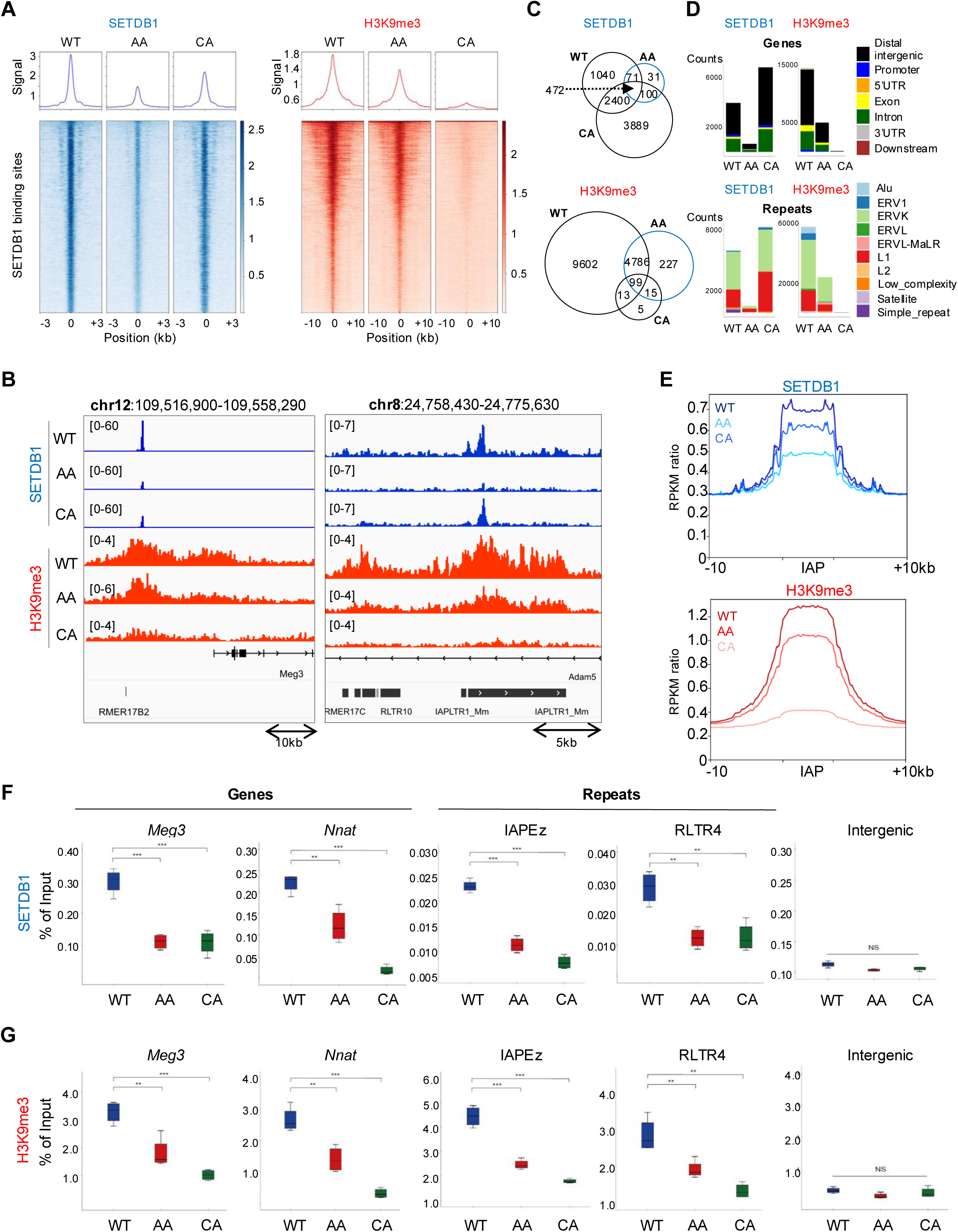
SETDB1 auto-methylation modulates its genomic binding capacity and influences the deposition of H3K9me3 at both coding genes and transposable elements. ChIP-seq and ChIP-qPCR data from *Setdb1* cKO mESCs re-expressing wild-type (WT) SETDB1, automethylation-deficient (AA) or catalytic-dead (CA) SETDB1 mutants. **A.** Heatmaps showing ChIP-seq signal intensity for Flag-SETDB1 and H3K9me3 across Flag-SETDB1-enriched regions, centered on peak summits and extended ±3 kb and ±10 kb, respectively. **B.** IGV genome browser tracks showing ChIP-seq signal for Flag-SETDB1 and H3K9me3 at representative loci—including the *Meg3* gene on chromosome 12 (chr12) and IAPLTR1_Mm repeat—on chromosome 8 (chr8) in mESCs expressing wild-type (WT), automethylation-deficient (AA), or catalytically inactive (CA) SETDB1 forms. **C.** Venn diagram representation of overlapping genomic regions occupied by Flag-SETDB1 SETDB1 in mouse embryonic stem cells (mESCs) expressing either the wild-type (WT), auto-methylation-deficient (AA), or catalytically inactive (CA) SETDB1 mutants (upper panel), and regions marked by H3K9me3 in the same cell lines (lower panel). **D.** Genomic annotation of Flag-SETDB1 binding sites across gene features and repeated families, based on ChIP-seq enrichment for Flag-SETDB1 and H3K9me3. Data shown represent enriched genomic regions consistently identified in both biological replicates. **E.** Pile-ups of the Flag-SETDB1 and H3K9me3 enrichment at IAP elements in *Setdb1* cKO mESCs reconstituted with wild-type (WT) SETDB1, auto-methylation-deficient (AA) or catalytic-dead (CA) SETDB1 forms. **F-G** Crosslinked ChIP-qPCR analysis of Flag-tagged SETDB1 (F) and H3K9me3 (G) at selected genes and repeated elements. An intergenic region lacking SETDB1 binding and H3K9me3 enrichment was used as a negative control. Statistical significance was assessed using Student’s *t*-test for normally distributed data, or Mann–Whitney’s unpaired test otherwise (n = 4 biological replicates). **p < 0.01; ***p < 0.001; NS: not significant.

Strikingly, although the SETDB1 automethylation-deficient AA mutant is less recruited to chromatin than the catalytic-dead CA mutant (**Figure 3A** left, **3B**), it still leads to significant H3K9me3 levels in contrast with the near-complete absence of H3K9me3 observed with the CA mutant (**Figure 3A** right, and **3B**). Notably, the H3K9me3 enrichment triggered by the SETDB1-AA mutant was weaker than that observed with SETDB1-WT. However, the associated H3K9me3 domains were broader in cells expressing SETDB1-AA form (**Figure 3A** right, **3B**, and **S3A**). This suggests defects in either the recruitment and/or chromatin-binding stability of the SETDB1-AA mutant.

The automethylation-deficient SETDB1 AA form shares 81% of its binding sites (543/674) with WT SETDB1 and 85% with the CA catalytic-dead mutant (572/674), while the CA catalytic-dead shares only 42% of its binding sites with WT SETDB1 (2872/6861) (**Figure 3C** upper). This might be explained by the overexpression of the CA mutant compared to WT and/or by the critical role of enzymatic activity in ensuring reliable targeting of SETDB1. As for H3K9me3 enrichments, the mESCs expressing the automethylation-deficient SETDB1-AA share 95% of H3K9me3-enriched regions (4885/5127) with mESCs expressing SETDB1-WT and 86% with the CA catalytic-dead mutant (114/132), while the catalytic-dead CA mutant shares 85% of its H3K9me3-enriched regions with SETDB1-WT (112/132) (**Figure 3C** lower).

Analysis of SETDB1 binding revealed approximately the same distribution across all forms, with approximately 60–70% of binding sites located in intergenic regions, around 25% in introns, and roughly 5% at promoters (**Figure 3D** upper). The majority of H3K9me3 enrichment in mESCs expressing WT SETDB1 and the AA automethylation-deficient mutant of SETDB1 is located in intergenic (∼60-65%) and intronic regions (∼20%) (**Figure 3D** upper). The distribution in coding sequences (<10%) or regulatory elements, such as promoters (<5%), showed less H3K9me3 enrichment.

In terms of chromatin states (**Figure S3B**), WT SETDB1 is enriched at intergenic regions (58%) and heterochromatin (20%), the catalytic-dead mutant shows globally the same distribution with more enrichment at intergenic regions (69%), while the AA automethylation-deficient mutant seems reduced in heterochromatin (17%). The H3K9me3 enrichment is barely detected in mESCs expressing the CA catalytic-dead SETDB1 mutant, and its distribution is reduced in all compartments in mESCs expressing the AA automethylation-deficient mutant, especially in intergenic and heterochromatin regions (**Figure S3B**).

Next, we asked whether the integrity of K1186/K1194 is required for SETDB1 recruitment to TEs, such as endogenous retroviral (ERV) and long interspersed nuclear element (LINE) elements, since SETDB1-deposited H3K9me3 is necessary for silencing of several ERV subfamilies in mESCs (Karimi et al., 2011; Matsui et al., 2010). A reduction of SETDB1 enrichment was apparent at IAP and more globally at ERV elements in mESCs expressing the AA automethylation-deficient SETDB1 (**Figures 3E** upper and **3F**, **Figure S3C**, **S3D** and **S3E**). We also checked whether the expression of the catalytically inactive SETDB1 affects its recruitment to ERVs and LINE-1 sequences. As it has been reported previously for the IAP elements (Karimi et al., 2011; Matsui et al., 2010), the inactivity of the enzyme only slightly affected SETDB1 enrichment levels at ERVs or LINE-1 sequences (**Figures 3E** and **3F**, **Figure S3C**, **S3D** and **S3E**).

Similar to the differences detected for H3K9me3 enrichment at genes in the presence of the automethylation-deficient and catalytically inactive mutants of SETDB1, the global H3K9me3 enrichment on TEs was barely detectable in mESCs expressing the catalytically inactive SETDB1 (CA), and it was reduced in those expressing the automethylation-deficient AA mutant, compared to the cells expressing WT SETDB1 (**Figures 3E** lower, **3G**, **S3D** and **S3F**).

We have confirmed by ChIP-qPCR that the automethylation-deficient SETDB1-AA is less recruited to target genes and TEs. We have tested a number of known SETDB1 target genes including *Meg3*, *Mest*, *Nnat* and *H19*, and TEs, including the ERV-K family repeats IAPEz, ETnERV and MMERVK10c, and the ERV-1 member RLTR4 (**Figure 3F** and **S3E**), and confirmed reduced binding of the automethylation-deficient SETDB1-AA mutant with a concomitant decrease in H3K9me3 enrichment at the promoter region of these genes (**Figure 3G** and **S3F**). These findings indicate that the integrity of K1186/K1194 in SETDB1 is required to maintain proviral silencing *via* efficient deposition of H3K9me3.

Altogether, these findings suggest that while the presence of lysines K1186 and K1194 affects SETDB1’s chromatin binding, its catalytic activity appears dispensable for this recruitment. Moreover, the presence of K1186/K1194 is essential for efficient SETDB1-mediated H3K9me3 deposition, underscoring their functional importance in epigenetic regulation.

### Integrity of SETDB1 K1186/K1194 is required for gene expression regulation and proviral silencing

To determine whether the integrity of the two automethylated lysines K1186 and K1194 of SETDB1 can regulate SETDB1-dependent gene regulation, we performed bulk RNA-seq on three biological replicates of *Setdb1* cKO TT2 mESCs re-expressing either WT SETDB1, CA catalytic-dead, or the AA automethylation-deficient SETDB1 mutants.

Multidimensional scaling (MDS) analysis revealed a clear separation of samples into three distinct clusters, corresponding to their respective biological replicates (**Figure 4A**). Overall, we found a significant number of differentially expressed genes in mESCs expressing SETDB1 mutant forms compared to those expressing the WT form (**Figure S4A**). Gene expression analysis of RNA-seq data, using thresholds of log₂ fold change > 0.5 and FDR < 0.05, identified 958 differentially expressed genes (DEGs) in mESCs expressing the AA automethylation-deficient mutant of SETDB1 compared to those expressing WT SETDB1. Among these, 825 genes were upregulated and 133 were downregulated (**Figure 4B** and **S4A**). In comparison, we found 2586 DEGs (1625 up- and 961 down-regulated) in mESCs expressing the CA catalytic-dead mutant compared to mESCs expressing WT SETDB1 (**Figures 4C** and **S4A**). Strikingly, 860 genes (740 upregulated and 120 downregulated) were commonly deregulated in mESCs expressing both the automethylation-deficient and catalytically-dead mutant of SETDB1 (**Figure 4D** and **S4A**). Notably, while the number of upregulated genes in the catalytically inactive CA mutant expressing cells was approximately twice that observed in the automethylation-deficient AA mutant, the number of downregulated genes exceeded an eightfold difference. This suggests that the automethylation-deficient AA mutant exerts a more limited effect on transcriptional repression compared to the catalytically inactive CA mutant.

**Figure 4.**
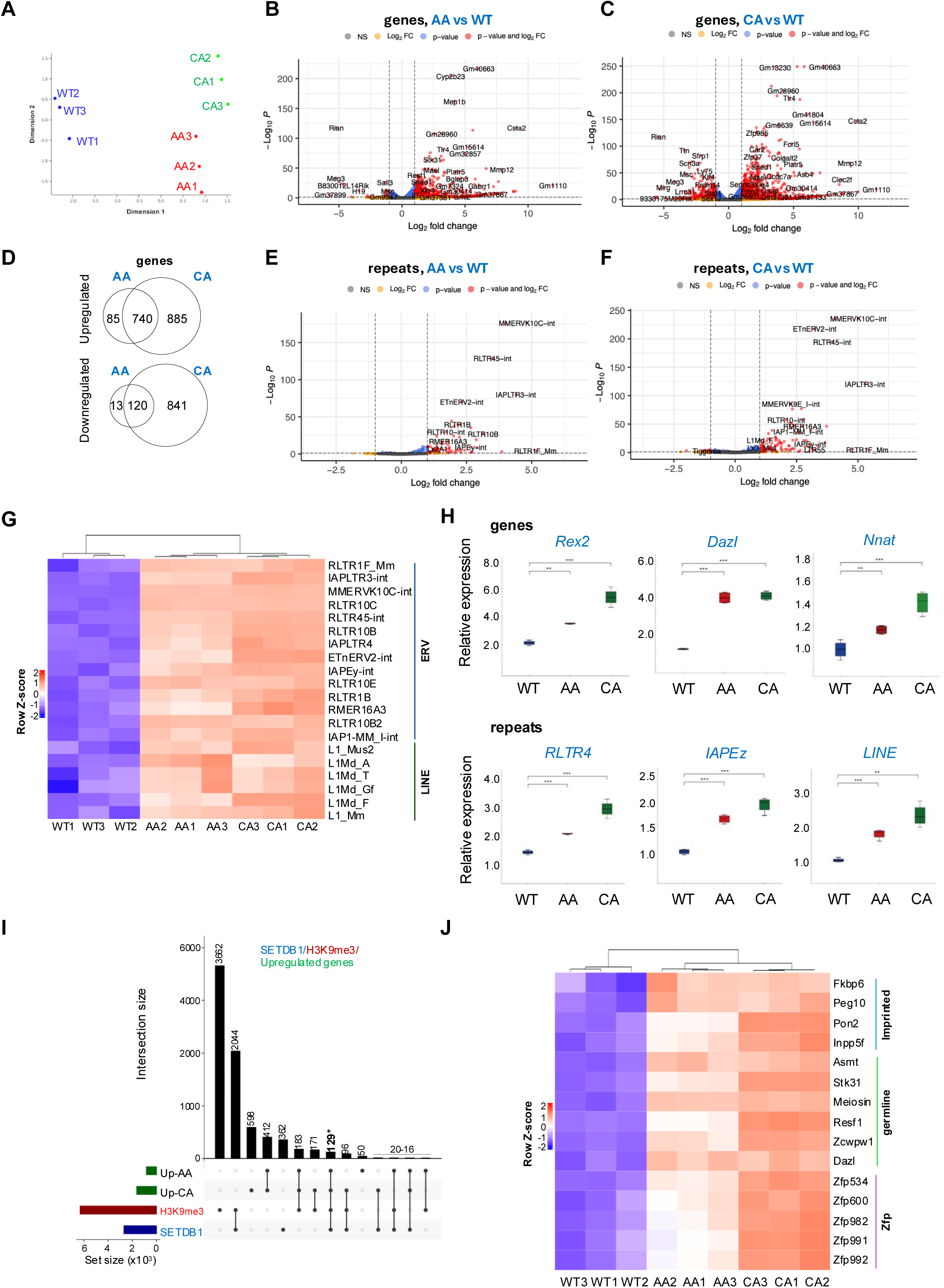
SETDB1 auto-methylation is required for its gene and repeat silencing activity. Total RNA-seq from and RT-qPCR data from *Setdb1* cKO mESCs re-expressing wild-type (WT) SETDB1, automethylation-deficient (AA) or catalytic-dead (CA) SETDB1 mutants. **A.** Multidimensional Scaling (MDS) plot of RNA-seq replicates (n = 3 independent experiments), showing clustering of samples (WT, AA, CA) based on global transcriptomic profiles. **B-C.** Volcano plot displaying differentially expressed genes between AA and WT-(**B**) and CA and WT-expressing cells (**C**), as measured by RNA-seq. Genes with an absolute log₂ fold change > 0.5 and a false discovery rate (FDR) < 0.05 are considered significantly regulated. Upregulated and downregulated genes meeting these criteria are highlighted in red. **D.** Venn diagram illustrating the overlap of differentially expressed genes (both upregulated and downregulated compared to WT) between AA and CA SETDB1-expressing cells, based on RNA-seq analysis. **E-F.** Volcano plot displaying differentially expressed repeated elements between AA and WT-(**E**) and CA and WT-expressing cells (**F**), as measured by RNA-seq. Repeated elements with an absolute log₂ fold change > 0.5 and a false discovery rate (FDR) < 0.05 are considered significantly regulated. Upregulated and downregulated repeats meeting these criteria are highlighted in red. **G.** Heatmap of Z-scored expression values for a representative set of differentially expressed repeated elements (ERVs and LINEs) across WT, AA, and CA mESCs biological replicates. Hierarchical clustering of samples highlights robust reproducibility across replicates. **H.** RT-qPCR analysis of selected deregulated genes (upper panel) and repeated elements (lower panel) in the indicated cell types (WT, AA, CA). mRNA levels were normalized to Cyclophilin A expression. Statistical significance was assessed using Student’s t-test for data with normal distribution, and Mann–Whitney’s unpaired test for non-parametric data (n = 4 biological replicates). **p < 0.01; ***p < 0.001 **I.** UpSet plot illustrating the intersection of Flag-SETDB1 and H3K9me3 ChIP-seq peaks with upregulated genes in AA and CA SETDB1-expressing mESCs. The plot quantifies the degree of overlap among datasets, with 129 shared genes highlighted in bold and marked with an asterisk. **J.** Heatmap of Z-scored expression values for a representative set of differentially expressed genes, including imprinted, germline-specific, and Zfp genes, across WT, AA, and CA mESC biological replicates. Hierarchical clustering of both genes and samples reveals distinct expression patterns and underscores the reproducibility across replicates.

Gene ontology (GO) analysis revealed that deregulated genes in mESCs expressing the AA automethylation-deficient SETDB1 mutant were mostly enriched for meiosis-related (or germline-specific genes), among others (**Figure S4B**). Interestingly, the GO categories enriched in mESCs expressing the CA catalytic-dead form of SETDB1 differed slightly from those in cells expressing the AA automethylation-deficient mutant (**Figure S4B** and **S4C**). This suggests that loss of catalytic activity has a broader impact on transcriptional regulation than the absence of automethylation at lysines K1186 and K1194.

Since SETDB1 is required for H3K9me3-mediated silencing of repeated elements in mESCs (Matsui et al., 2010), we investigated whether the presence of the two automethylated lysines, K1186 and K1194 is necessary for SETDB1-dependent transcriptional silencing of ERV and LINE sequences in mESCs. The RNAseq data, confirmed by RT-qPCR for few targets, showed that expression of the automethylation-deficient AA mutant of SETDB1 in mESCs resulted in de-repression of many ERV and LINE elements (**Figures 4E**, **4G** and **4H** lower, **Figure S4D**) in a comparable way as the expression of the CA catalytic-dead SETDB1 mutant (**Figure 4F** and **4G, 4H** lower**, S4D**).

Among the upregulated genes in cells expressing the AA and CA SETDB1 mutants, we identified those that are normally occupied by SETDB1 and enriched in H3K9me3 in WT SETDB1-expressing mESCs. This analysis revealed 129 shared genes (**Figures 4I** and **4J**), including germline genes, imprinted genes, and members of the zinc finger protein (Zfp) family, all known to be directly repressed by SETDB1 through H3K9me3 and DNA methylation in mESCs (Karimi et al., 2011). We have confirmed some of the RNAseq results by RT-qPCR, including for *Rex2*, *Dazl*, *Mest*, *Stk31*, *Tuba3a*, and *Hormad1* expression (**Figures 4H** upper**, 4J**, and **Figure S4D**). These findings demonstrate that disruption of SETDB1, whether through loss of catalytic activity or automethylation at K1186/K1194, is sufficient to trigger transcriptional reactivation of its target genes.

Altogether, these findings reveal that the expression of the automethylation-deficient SETDB1 mutant disrupts SETDB1/H3K9me3-mediated silencing of coding genes and many ERVs in mESCs, and its effects resemble those observed in cells expressing the catalytically inactive form of SETDB1, albeit to a lesser extent.

These results suggest that expression of the SETDB1 automethylation-deficient mutant in mESCs could lead to a loss of pluripotency, an event that has been demonstrated in mESCs after loss of SETDB1 (Dodge et al., 2004; Yeap et al., 2009).

### SETDB1 automethylation affects its protein-protein interactions

The methyltransferase activity of SETDB1 is not directly affected by the double K1186/K1194 mutation (**Figure 2A**), yet the H3K9me3 levels are perturbed in mESCs expressing the automethylation-deficient AA mutant (**Figure 3** and **S3**), suggesting deregulated H3K9me3 deposition and/or spreading mechanisms. Thus, we postulated that SETDB1 automethylation on K1186 and K1194 is involved in modulating other features, such as key protein–protein interactions, especially with methyl-lysine readers cooperating with SETDB1 in the establishment and/or spreading of H3K9me3.

To address this, we analyzed the immunoprecipitated Myc-tagged products from *Setdb1* knockout MEFs nuclear extracts by WB (as in **Figure S1E** and **2A**). Interestingly, both the integrity of K1186/K1194 and SETDB1 catalytic activity were required for its interaction with several heterochromatin-inducing complexes, such as the chromodomain-containing proteins Suv39h1, HP1γ, and CDYL (Chromodomain Y-Like) (**Figure 5A**).

**Figure 5.**
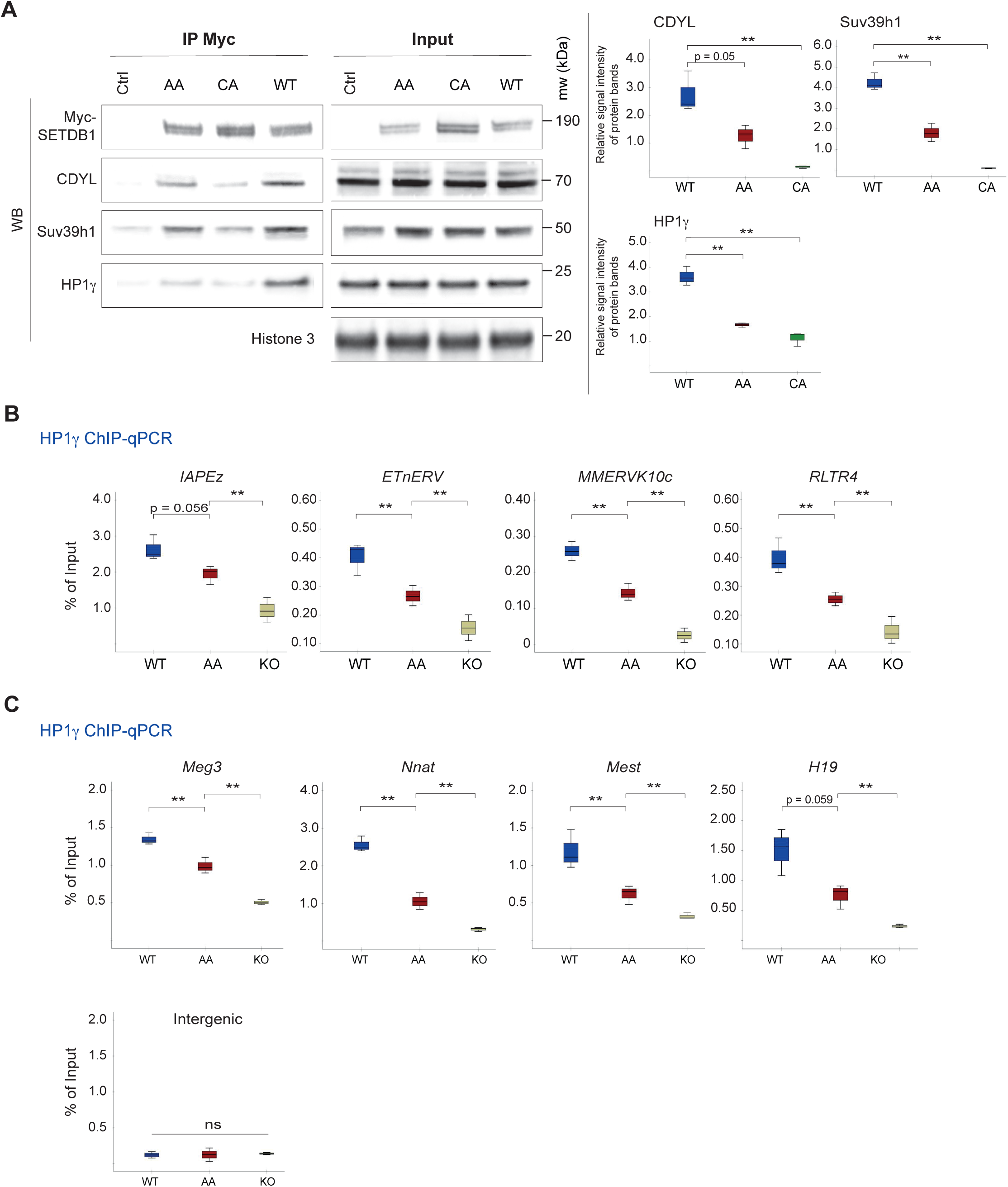
Methylation status of SETDB1 affects its interaction with the chromodomain-containing proteins and its cooperative role with HP1γ in H3K9me3 establishment and spreading. **A.** Decrease of interaction between automethylation-deficient mutant (AA) and catalytically inactive (CA) SETDB1 with the chromodomain-containing proteins Suv39h1, CDYL, and HP1γ. Myc-tagged full-length wild-type (WT), automethylation-deficient (AA) or catalytic-inactive (C1243A) SETDB1forms were transiently expressed in *Setdb1* KO MEFs. Nuclear extracts were used for immunoprecipitation (IP) with Myc-Trap technology. The resulting precipitates were then subjected to Western blot (WB) with the indicated antibodies (left panel). The results for the inputs are in the right panel. WB signals were quantified and presented as the ratio between the analyzed proteins over the Myc-tagged SETDB1 (n = 3 biological replicates) **: p < 0.05 (right panel). **B-C.** Crosslinked ChIP-qPCR analysis of HP1γ in *Setdb1* cKO mESCs and cells re-expressing WT or AA mutant SETDB1 after 96h of tamoxifen treatment, Repeated elements (B) and imprinted genes (C). Statistical significance was assessed using Student’s *t*-test for normally distributed data, or Mann–Whitney’s unpaired test otherwise (n = 4 biological replicates). **p < 0.01; NS: not significant.

Given that HP1, a key heterochromatin component, recognizes methylated lysines and is essential for proviral silencing (Bulut-Karslioglu et al., 2014), we hypothesized that SETDB1 automethylation could modulate HP1 recruitment to chromatin. Hence, we asked whether HP1γ recruitment to chromatin is dependent on the integrity of K1186/K1194 of SETDB1. To test this, we performed ChIP-qPCR assays and found that HP1γ recruitment to many known SETDB1 target genes (*Nnat*, *Mest*, *Meg3* and *H19*) or repeats (including the ERV-K family repeats IAPEz, ETnERV and MMERK10c, and the ERV-1 member RLTR4) is impaired when lysines K1186 and K1194 are mutated (**Figures 5B** and **5C**). Notably, since the expression of the automethylation-deficient AA SETDB1 mutant results in reduced enrichment of H3K9me3, the reduced HP1γ recruitment could be attributed to both reduced levels of H3K9me3 and its reduced binding to automethylation-deficient AA SETDB1.

In summary, our findings show that SETDB1 automethylation on K1186 and K1194 regulates its interaction with methyl-lysine reader proteins such as HP1γ, and at least partially regulates its recruitment to chromatin.

## DISCUSSION

SETDB1, a key histone lysine methyltransferase involved in transcriptional repression, is extensively decorated by PTMs that influence its enzymatic activity, stability, and protein-protein interactions (https://Phosphosite.org). Our findings demonstrate that SETDB1 undergoes automethylation at lysines K1170 and K1178 within its catalytic SET domain, modifications that do not affect SETDB1’s enzymatic activity but are essential for its interaction with heterochromatin-associated proteins such as HP1γ, Suv39h1, and CDYL. Mutation of these residues disrupted methylation and compromised binding, indicating that both lysines are functionally indispensable (**Figure S5**).

These lysines lie within sequence motifs resembling the conserved ARKS motif of histone H3 tail, suggesting a histone-mimicry mechanism in which SETDB1 recruits chromodomain-containing proteins in a manner analogous to histone H3K9me3. This mimicry may extend heterochromatin regulation to non-histone proteins, and represents a critical element of SETDB1’s function. Notably, a recent study identified two other histone mimic methylated sites of SETDB1 K489/K490 and K1162, which are important for the HP1α binding to SETDB1 (https://doi.org/10.1101/2025.01.21.634156). These findings suggest that histone-mimic automethylation may be a broader regulatory theme, extending beyond K1170 and K1178 identified in this study, and K489/K490 and K1162, and providing multiple “docking” sites for reader proteins. Moreover, at least one adjacent serine residue (S1179, https://Phosphosite.org) was found to be phosphorylated, potentially establishing a phospho/methyl switch similar to the H3K9me3/S10ph mechanism (Fischle et al., 2005), further increasing the regulatory complexity of SETDB1 activity by adding another layer of regulation: stabilizing or disrupting reader protein binding depending on phosphorylation state (e.g., via kinases acting in response to signaling events such as DNA damage).

In addition to methylation, SETDB1 is also ubiquitinated at K867 (Ishimoto et al., 2016), a modification required for its catalytic activity and regulation of target gene expression, highlighting the integration of multiple PTMs in fine-tuning SETDB1 function.

Our ChIP-seq analysis reveals that lysines K1186 and K1194 play a key role in stabilizing SETDB1 at its genomic targets, which in turn affects the efficiency of H3K9me3 establishment. The observation that the catalytic-dead mutant remains detectable at target loci suggests that initial recruitment is largely independent of enzymatic activity. By contrast, the weaker enrichment of the automethylation-deficient AA mutant when compared to both WT and catalytic-dead CA mutant, despite its unperturbed enzymatic activity, highlights the importance of these residues in sustaining SETDB1 binding. Importantly, SETDB1 methylates H3K9 in a distributive manner, adding one methyl group per binding event and dissociating between rounds (Basavapathruni et al., 2016), suggesting that stable chromatin association is essential to achieve full trimethylation. Interactions with chromodomain-containing proteins may therefore provide an additional stabilizing mechanism, ensuring sufficient residence time for complete H3K9me3 deposition. This model aligns with proposed functions of histone-mimic motifs in heterochromatin spreading (https://doi.org/10.1101/2025.01.21.634156), but also raises the possibility that similar stabilization mechanisms could be broadly employed to regulate the fidelity of other histone-modifying enzymes.

Failure to stabilize binding and properly establish H3K9me3 at target sites results in partial de-repression of SETDB1-targeted genes and repetitive elements, particularly ERVs, in the presence of the automethylation-deficient AA mutant. Although there was partial establishment of H3K9me3, it was insufficient for maintaining transcriptional repression of a number of targeted sequences, including germline, imprinted, and *Znf* family genes, and several families of ERVs. Importantly, the extent of transcriptional changes caused by the AA mutant is milder compared to the catalytically inactive CA mutant, which induces a broader spectrum of gene deregulation. This suggests that the catalytic activity of SETDB1 remains the dominant determinant of repression, while automethylation at lysines K1186/K1194 contributes to the stability and efficiency of SETDB1-mediated silencing, and its disruption is sufficient to compromise transcriptional control. Notably, while some loci exhibited lower enrichment of H3K9me3 in the presence of automethylation-deficient AA SETDB1 mutant, they were not derepressed, suggesting that compensatory mechanisms may be at play. However, these mechanisms appear insufficient to fully restore the expression pattern, underscoring the central role of SETDB1 in maintaining heterochromatin integrity in mESCs, especially at ERVs.

HP1 proteins recognize methylated lysines to promote heterochromatin formation and proviral silencing (Bulut-Karslioglu et al., 2014). Our results reveal a critical role for lysines K1186 and K1196 in SETDB1-mediated recruitment of HP1γ to chromatin. ChIP-qPCR analyses showed that mutating K1186/K1196 significantly impairs HP1γ recruitment to several SETDB1 target genes and repeated elements. Based on our data, we cannot distinguish whether HP1γ recruitment is impaired in the presence of the automethylation-deficient AA mutant due to the loss of the recognition site for HP1 or its associated cofactors on SETDB1 itself or due to the partial reduction of H3K9me3 establishment. Nevertheless, our findings identify K1186 and K1194 as critical residues for HP1γ recruitment. Our data, in agreement with a recent study (https://doi.org/10.1101/2025.01.21.634156), further suggest that SETDB1’s interaction with repressor complexes (e.g., CDYL) may be modulated by SETDB1 methylation state, potentially integrating histone mimicry with repressor complex recruitment, highlighting a previously unappreciated layer of regulation in SETDB1-mediated heterochromatin formation.

Finally, our previous work showed that SETDB1 forms a megacomplex with SUV39H1 and other H3K9 KMTs (Fritsch et al., 2010), which participate in the maintenance of heterochromatin. Our current observation of the role of automethylation in the regulation of the SETDB1/Suv39h1 interaction provides a mechanistic means to understand the regulation of such megacomplexes.

In conclusion, our study reveals that SETDB1 auto-methylation at two histone-mimic motifs (K1170/K1178 in human and K1186/K1194 in mouse) is critical for its function and ability to engage chromatin regulators required for heterochromatin formation (**Figure S5**). These PTMs, along with ubiquitination and potential phosphorylation, form a regulatory hub that modulates SETDB1’s role in repressing ERVs, imprinted genes, and repetitive elements. The similarity of SETDB1’s methylated motifs to histone sequences underscores the extension of chromatin regulatory logic to non-histone proteins. These findings highlight the broader significance of PTMs in safeguarding genome stability and orchestrating epigenetic control across diverse genomic contexts.

## Limitations of the study

We used K-to-A mutations to mimic the unmethylated form of SETDB1; however, lysines are subject to multiple PTMs beyond methylation, including acetylation and sumoylation. While we did not detect these modifications at K1186 and K1194 by mass spectrometry, we cannot formally exclude that loss of other potential PTMs at these sites contributes to the observed phenotypes. The absence of reliable methyl-mimetic lysine substitutions precludes direct demonstration that methylation *per se*, rather than lysine residues themselves, is critical for SETDB1 function.

The reduced HP1γ binding observed with K-to-A mutants could result from either direct loss of lysine-dependent interactions or indirect effects due to decreased H3K9me3 levels at SETDB1 target sites. Finally, our experiments were performed in ESCs cultured under serum + LIF conditions, where DNA methylation already marks a subset of repeat elements, potentially mitigating the full impact of H3K9me3 loss.

## RESOURCE AVAILABILITY

### Lead contact

Further information and request for resources and reagents should be directed to the lead contact, Dr Slimane Ait-Si-Ali.

### Materials availability

Materials generated in this study are available from the lead contact upon request.

### Data availability

- All NGS sequencing data that support the findings of this study are available under the accession numbers XXXXX and XXXXX.
- All data needed to evaluate the conclusions in the paper are present in the paper and/or the Supplementary Materials.
- Further details necessary for reanalysis of the data presented in this paper are available from the lead contact upon request.

## ACKNOWLEDGEMENTS

Work in the Ait-Si-Ali lab was supported by the Association Française contre les Myopathies Telethon (AFM-Telethon, grant # 22480, to S Ait-Si-Ali); Fondation pour la Recherche Médicale (FRM, « Equipe FRM » grant # DEQ20160334922, to S Ait-Si-Ali); Agence Nationale de la Recherche (ANR, grants ANR-17-CE12-0010-01 – MuSiC, ANR-22-CE14-0068-03 – EpiMuSe, and ANR-24-CE45-0690-03 - MultiPhysC2M to S Ait-Si-Ali), Université Paris Diderot (now Université Paris Cité) and the “Who Am I?” Laboratory of Excellence, # ANR-11-LABX-0071, to S Ait-Si-Ali, funded by the French Government through its “Investments for the Future” program, operated by the ANR under grant #ANR-11-IDEX-0005-01, and IDEX “Investissement d’Avenir” launched by the French Government and implemented by ANR, with the reference ANR-18-IdEx-0001 in the frame of Emergence call from Université Paris Cité. P.A-C.T was supported by PhD fellowships from the Administrative Department of Science, Technology and Innovation (COLCIENCIAS); Universidad del Rosario (Becas para Apoyo para Estudiantes Doctorales 2017); Colombian Institute of Educational Credit and Technical Studies Abroad (ICETEX); French Government Agency Campus France (Eiffel Excellence Scholarship Program); Fondation ARC pour la Recherche sur le Cancer; and a transition post-doc fellowship from the LABEX Who am I?. R.R. has been supported by a DIM Biotherapies – Paris and LABEX “Who am I?” (Université Paris Cité, ex Université Paris Diderot) fellowships.

We thank P Yoichi Shinkai for generous sharing of biological material. We thank the (EPI)2 Imaging platform (managed by Sandra Piquet) - UMR7216 Epigenetic and Cell Fate Center, for access to instruments and technical advice. We thank Dr Lauriane Fritsch for initiation of the project. We thank members of the Ait-Si-Ali lab and Epigenetics and Cell Fate department for helpful discussions during the group and department meetings and critical reading of the manuscript. The authors acknowledge the support of the Freiburg Galaxy Team: Person *X* and Björn Grüning, Bioinformatics, University of Freiburg (Germany), funded by the German Federal Ministry of Education and Research BMFTR grant 031 A538A de.NBI-RBC and the Ministry of Science, Research and the Arts Baden-Württemberg (MWK) within the framework of LIBIS/de.NBI Freiburg.

## AUTHOR CONTRIBUTIONS

Conceptualization, S.A.; Methodology Software; G.V.; Formal Analysis; P.A-C.T, E.B., L.D., G.V., R.R., G.C., V.C., V.J.; Investigation; P.A-C.T, E.B., L.D., G.V., R.R., Data Curation, P.A-C.T, E.B., L.D., G.V., R.R., G.C., V.C.; Writing – Original draft; S.A., Writing – Review & Editing; S.A., E.B., P.A-C.T, G.V.; Supervision, S.A.; Funding Acquisition, S.A.; Project Administration, S.A.

## MATERIAL AND METHODS

### RESOURCES TABLE

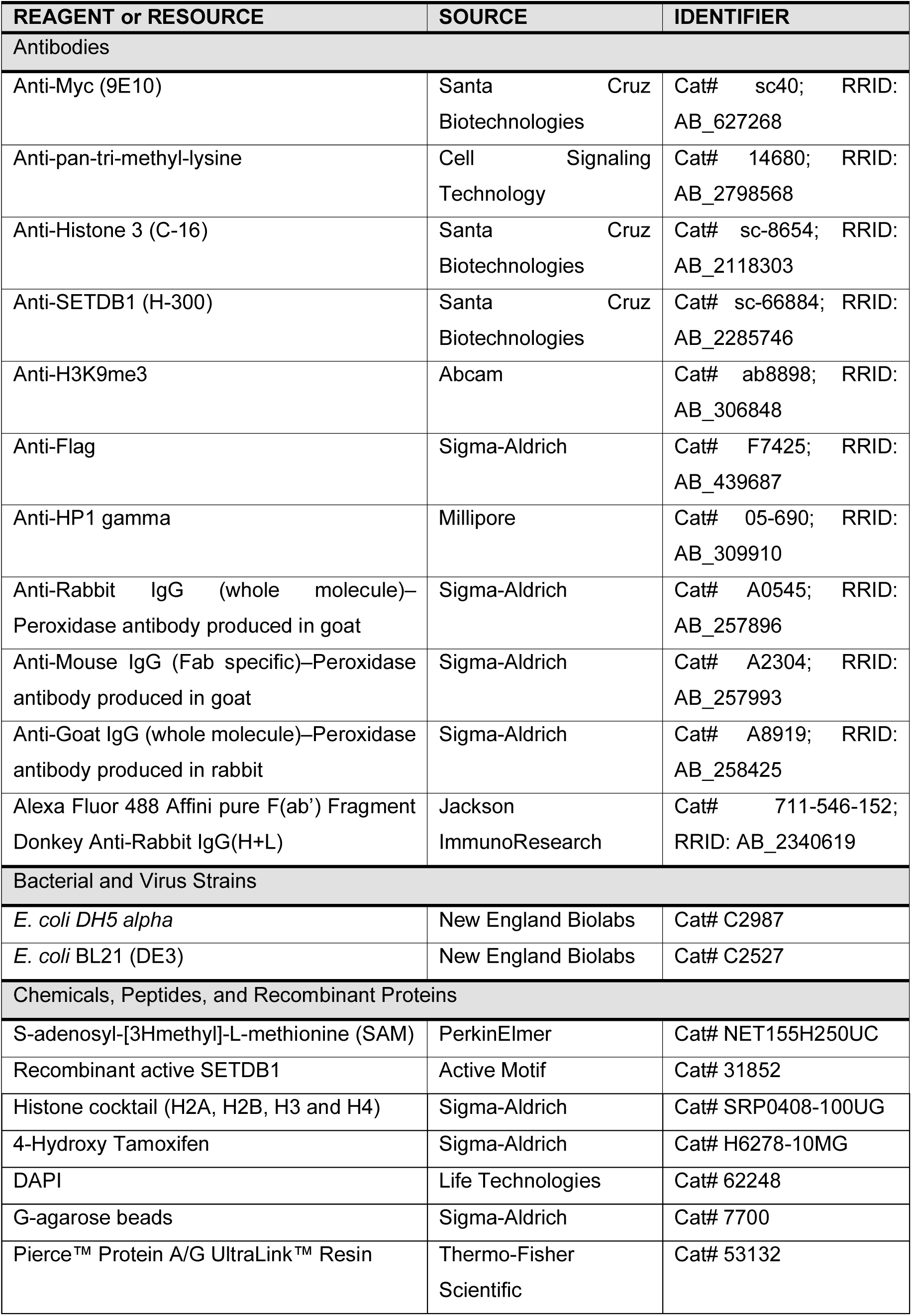
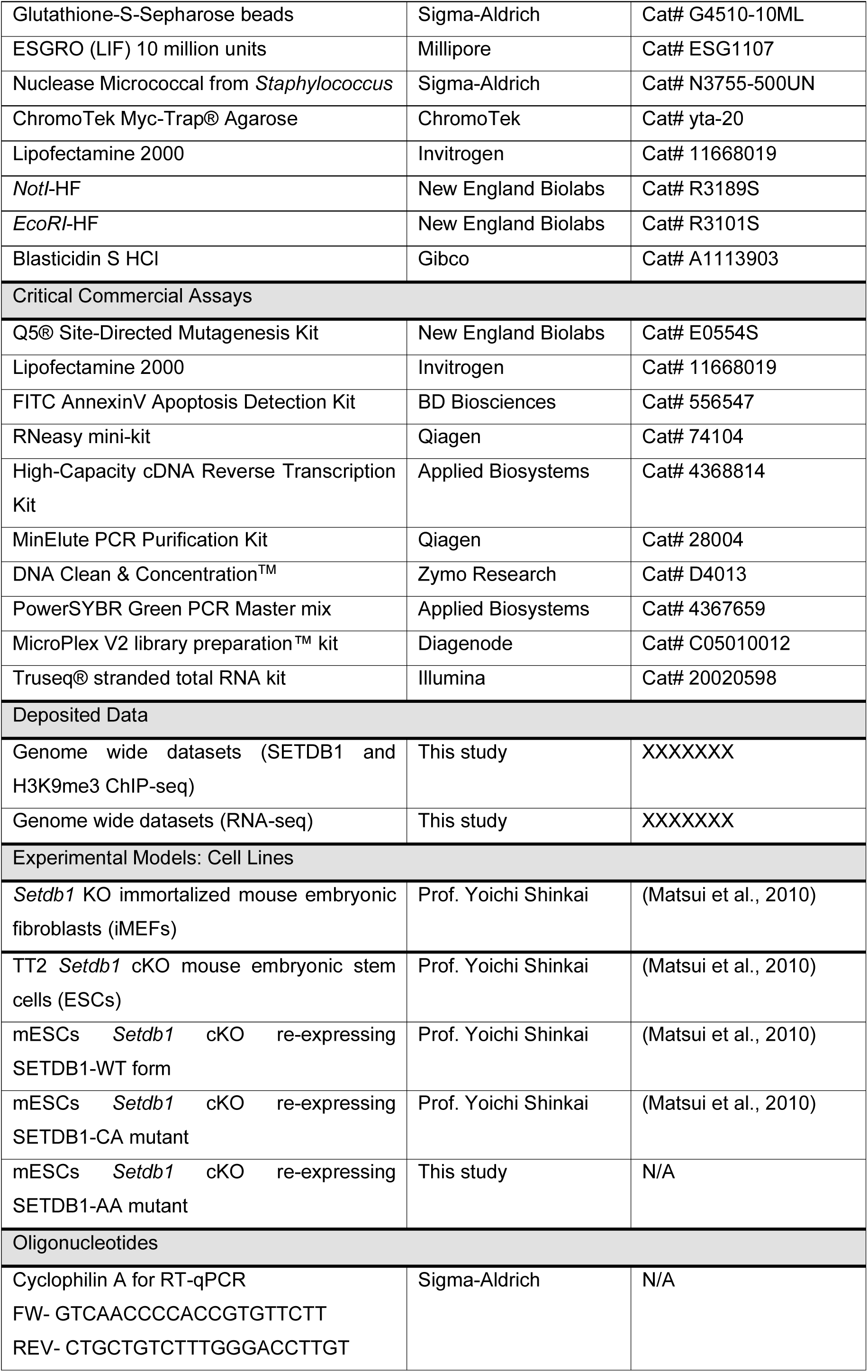
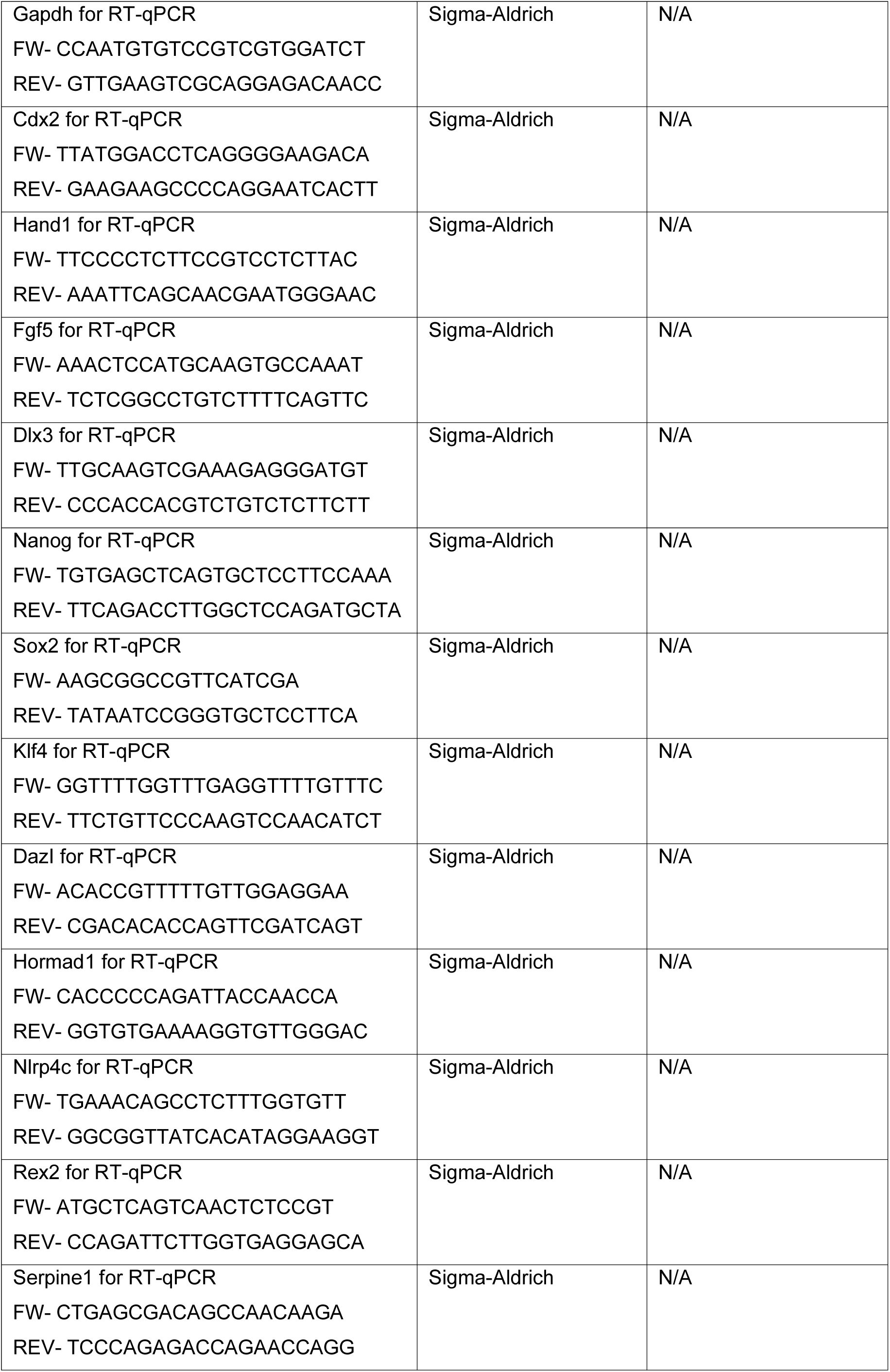
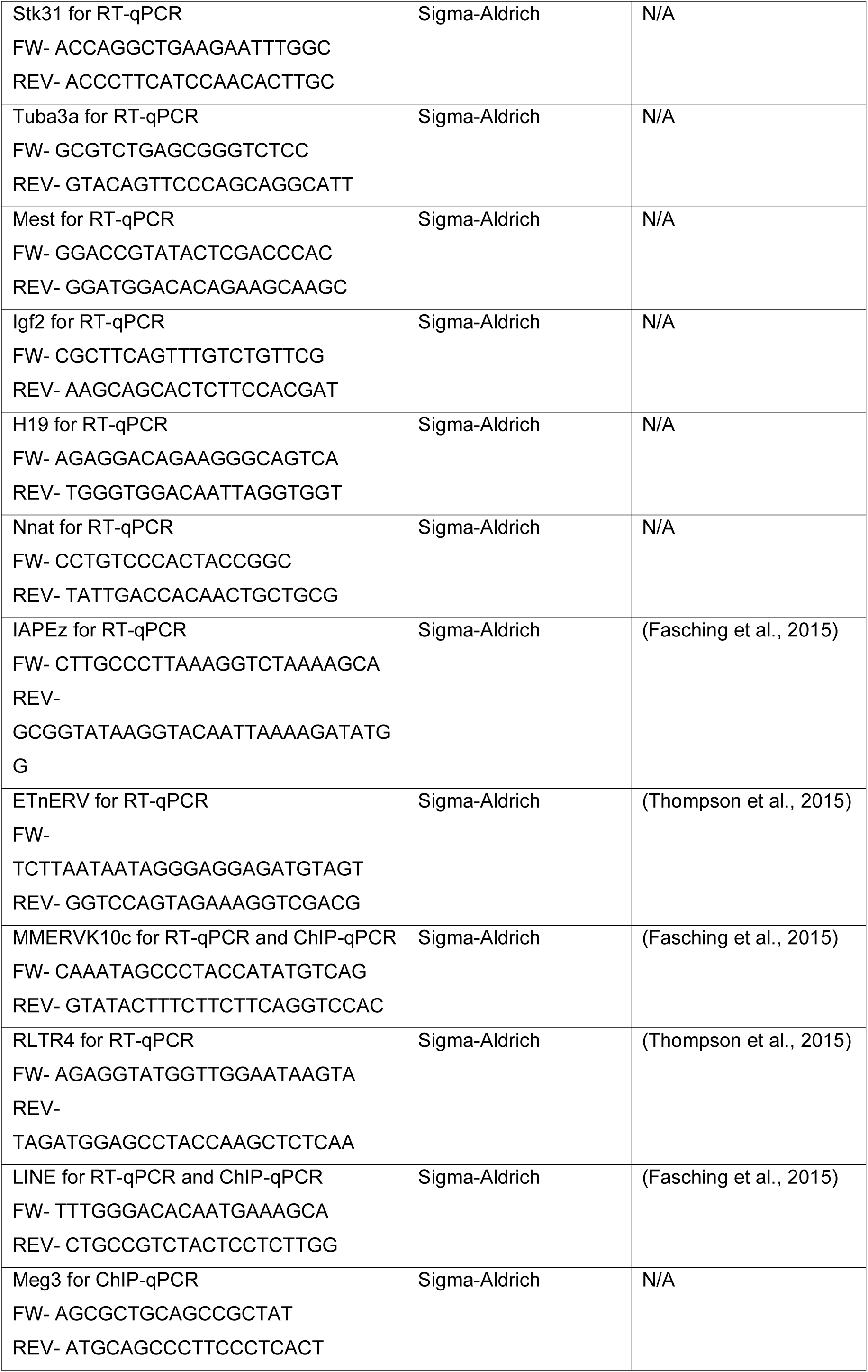
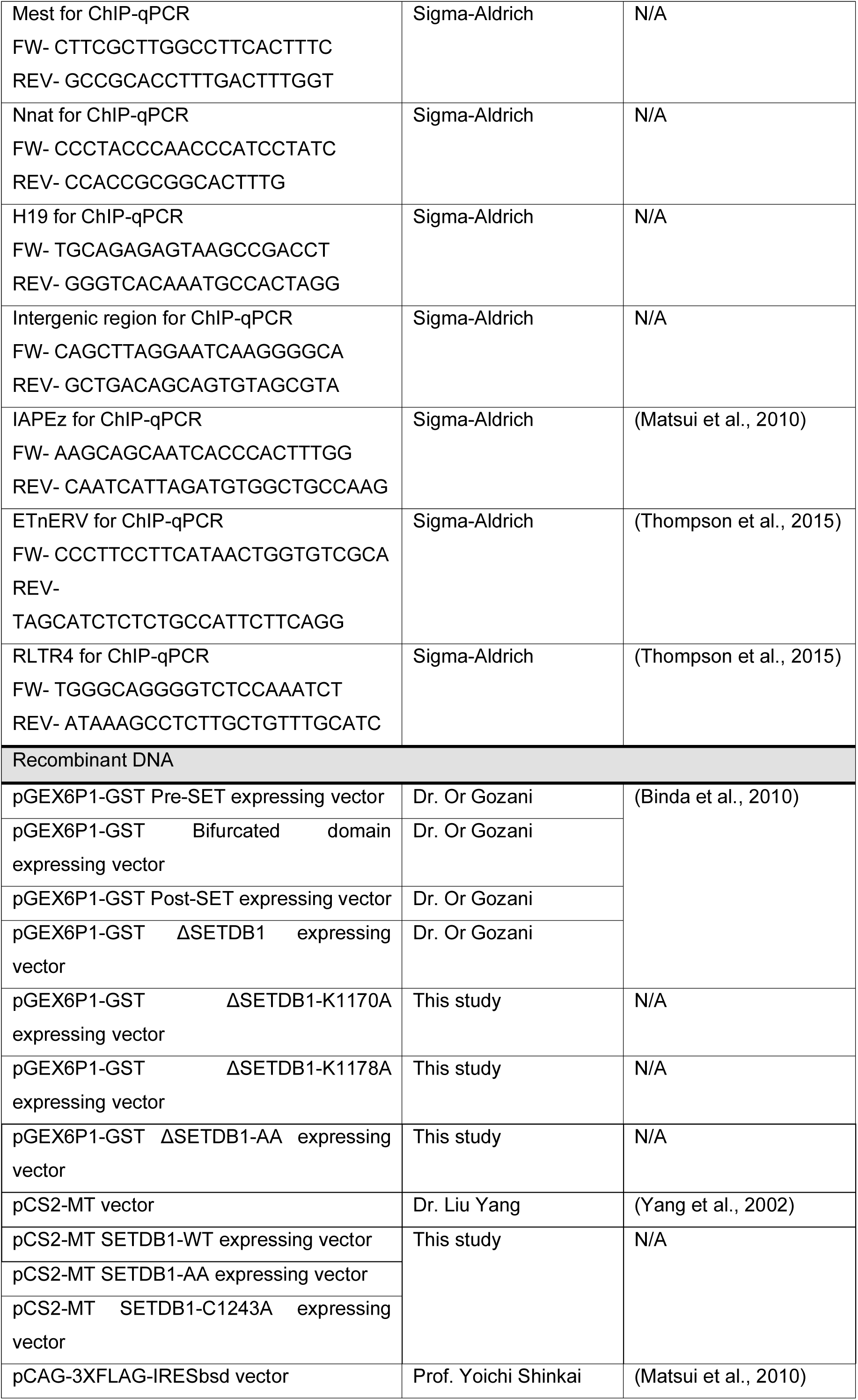
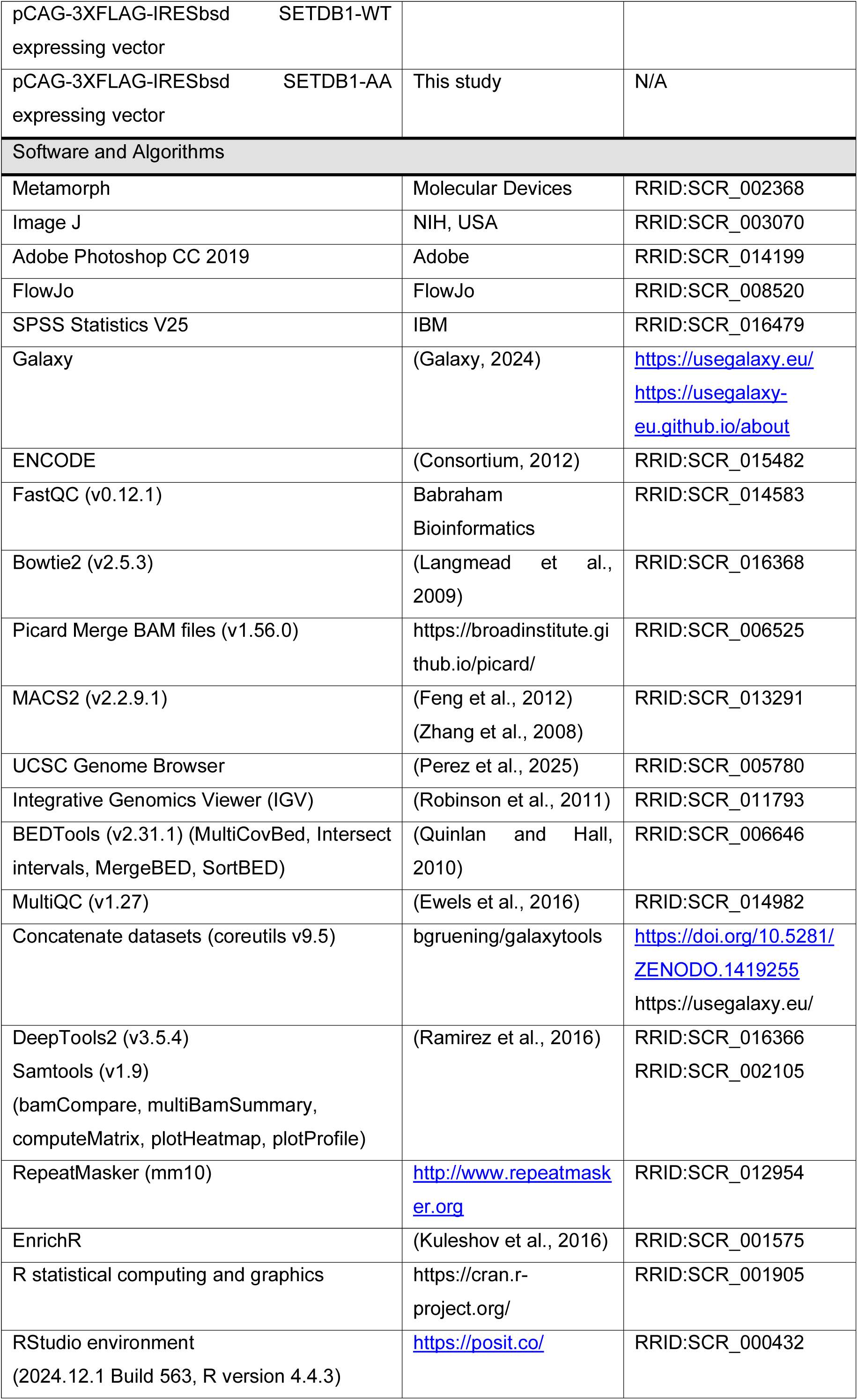
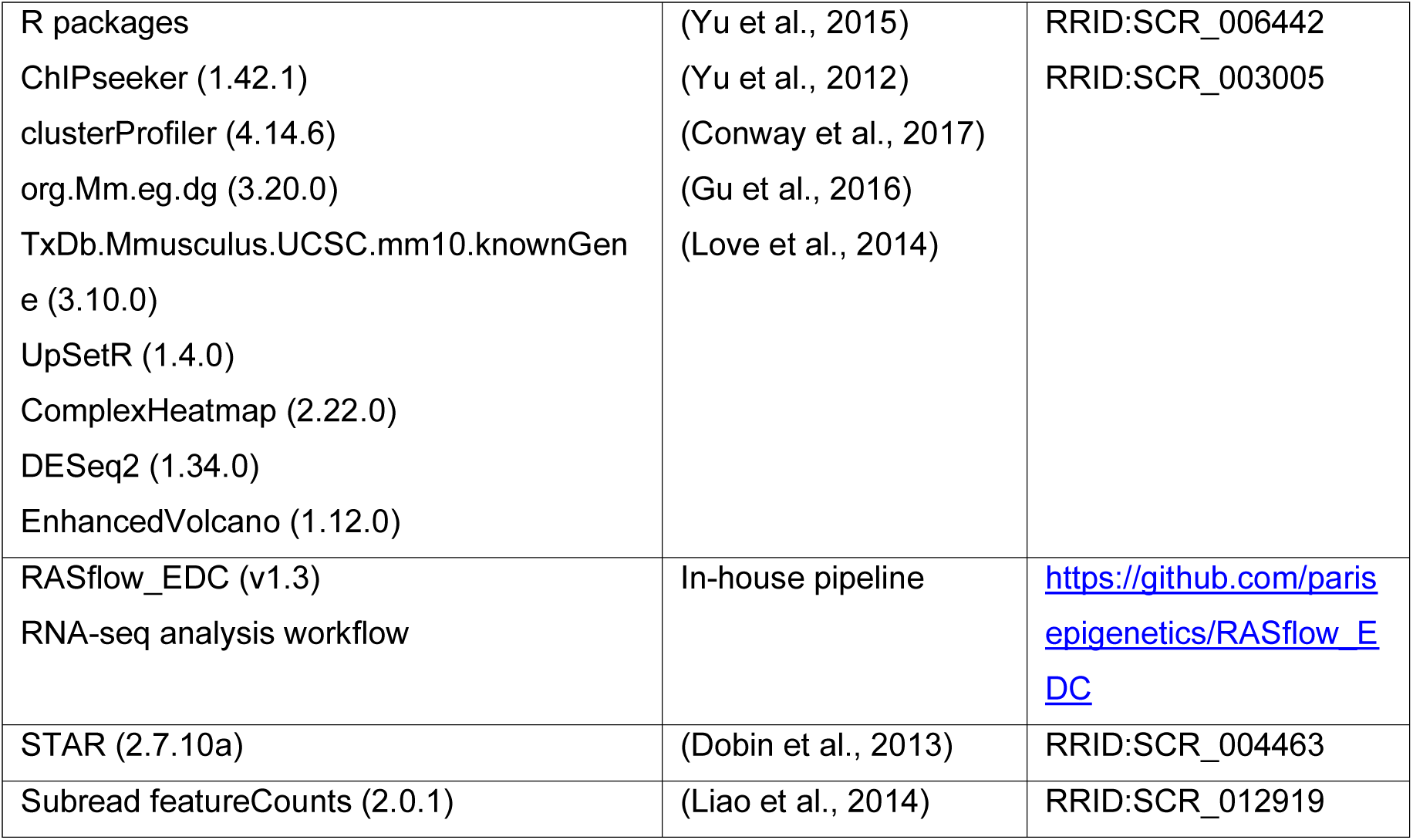

### EXPERIMENTAL MODEL AND SUBJECT DETAILS

#### Cell Culture and Cell Lines

*Setdb1* KO mouse embryonic fibroblasts (MEFs) and were kindly donated by Prof. Yoichi Shinkai and previously described (Matsui et al., 2010). *Setdb1* KO MEFs were cultured in DMEM (Sigma) supplemented with 15% fetal calf serum (Gibco) and 1% penicillin/streptomycin (Sigma). MEFs were maintained at 37°C and 5% CO_2_.

*Setdb1* cKO mouse embryonic stem cells (mESCs) were established by the group of Prof. Yoichi Shinkai *via* standard gene targeting procedures (Matsui et al., 2010). To generate the *Setdb1* cKO mESC line, Cre recombinase and estrogen receptor (Cre-ER) fusion gene was introduced into a clone containing targeted *Setdb1* cKO and KO alleles. We also used *Setdb1* cKO (re)-expressing 3XFlag-SETDB1 wild-type, as well as catalytically inactive SETDB1, kindly donated by Prof. Yoichi Shinkai (Matsui et al., 2010).

All mESCs were cultured in Dulbecco’s modified Eagle’s Medium DMEM (Sigma) supplemented with 15% fetal calf serum (Gibco), 1% penicillin/streptomycin (Sigma), 0.1 mM β-mercaptoethanol (Thermo), 1 mM nonessential amino acids (Sigma), and 1000 U/mL of Leukemia Inhibitory Factor (LIF) (Millipore). All mESC lines were cultured in standard feeder-free conditions with 0.2% gelatin, maintained at 37°C and 8% CO_2_. To induce deletion of the SETDB1 cKO allele, mESCs were cultured in 800nM 4-Hydroxy Tamoxifen (Sigma) for 4 days.

### METHOD DETAILS

#### Establishment of cell lines

Full-length SETDB1 cDNA was cloned into the *NotI* and *EcoRI* sites of the pCS2 vector donated by Dr. Liu Yang (Yang et al., 2002), which contains a Myc epitope tag. In order to generate the catalytic-dead (C1243A) SETDB1 mutant, or the double-lysine SETDB1 mutant (K1770A and K1778A), we used the Q5® Site-Directed Mutagenesis Kit (New England Biolabs) according to the manufacturer’s recommendations. Four different Myc-tagged SETDB1 cDNAs, including full-length (WT), automethylation-deficient (AA) and catalytic-inactive (C1243A, named CA) forms, as well as the empty pCS2 vector were transiently expressed in *Setdb1* KO MEFs *via* Lipofectamine 2000 (Invitrogen).

SETDB1 cDNA containing the mutations K1770A and K1778A was cloned into the *NotI* and *EcoRI* sites of the pCAG-3XFLAG-IRESbsd vector used by Prof. Yoichi Shinkai Lab (Matsui et al., 2010). To generate mESCs stably expressing the double-lysine SETDB1 mutant, the vector was introduced into *Setdb1* cKO mESCs *via* Lipofectamine 2000 (Invitrogen). Clones were selected in medium containing 7 μg/mL Blasticidin (Gibco).

#### Expression and purification of GST-tagged proteins

GST-tagged SETDB1 truncated mutants (pre-SET, Bifurcate domain, post-SET, and ΔSETDB1) were kindly donated by Prof. Or Gozani (Binda et al., 2010). The lysine mutations K1170A and/or K1178A were generated by site-directed mutagenesis as previously described. GST fusion proteins were transformed into *E. coli* Bl21 (DE3) (New England Biolabs). An overnight 20 mL culture from a single colony was grown at 37°C in LB medium containing 100 mg/mL of ampicillin. Then, the culture volume was increased to 500 mL. Gene expression was induced at OD=0.7 by adding 1 mM IPTG at 37°C for 2 h. After centrifugation, bacteria were resuspended in Lysis buffer (20 mM Tris pH 7.6, 150 mM NaCl, 0.5% Triton X-100, 0.01% NP40, 15 mM KCl, 10% Glycerol, 1 mM DTT, 0.5 mg/mL Lysozyme) and incubated for 30 min on ice. Then, the suspension was disrupted by sonication for 10 min (30 sec ON, 30 sec OFF) at high frequency (Bioruptor Diagenode). Bacterial debris were removed by centrifugation at 12000 rpm for 30 min at 4°C. GST-tagged proteins were bound to Glutathione-S-Sepharose beads (Sigma) for 2 h, at 4°C. After washing with Wash buffer (50 mM Tris pH 7.6, 300 mM NaCl, 0.5% Triton-X-100 and 1 mM DTT), bound proteins were eluted with Elution buffer (100 mM Tris pH 8.0, 15 mM Glutathione, 1 mM DTT), collected by centrifugation, and stored at -80°C. Proteins were resolved on 4%–12% NuPAGE gels (Invitrogen) in MES running buffer and visualized by SimplyBlue SafeStain (Invitrogen).

#### *In vitro* histone methyltransferase assay

GST-purified proteins (1 μg) or products from immunoprecipitation were incubated in a solution of reaction buffer (100 mM Tris pH 8, 40 mM KCl, 20 mM MgCl_2_, 20 mM B-Mercaptoethanol, 500 mM Sucrose), 30 mM S-adenosyl-[3Hmethyl]-L-methionine (SAM) and either 2 μg recombinant active SETDB1 (Active Motif) or 5 μg Histone cocktail (H2A, H2B, H3 and H4). The reactions were incubated for 2 h at 37°C. Then, the samples separated by SDS-PAGE, and the proteins were fixed with 50% methanol and 10% acetic acid and visualized by SimplyBlue SafeStain (Invitrogen). Finally, the gel was incubated with En3hance and exposed to autoradiography films overnight at -80°C.

#### Immunofluorescence (IF)

Cells were grown on glass coverslips, which had been coated with 10 mg/ml laminin for 1 h at 37 °C and rinsed with PBS. mESCs cells were fixed with 4% formaldehyde 20 min, incubated with 50 mM NH4Cl for 10 min to quench formaldehyde, and permeabilized with 0.2% Triton-X-100 for 10 min. Primary (SETDB1. Cat# sc-66884; Santa Cruz Biotechnologies) and secondary antibody (Alexa Fluor 488 Affini pure F(ab’) Fragment Donkey Anti-Rabbit IgG. Cat# 711-546-152; Jackson ImmunoResearch) were diluted in PBS containing 2% SVF serum and 0.1% Tween and incubated overnight at 4°C or 1 h at room temperature, respectively. DNA was stained with 1 μg/mL DAPI (Life Technologies). Coverslips were mounted with Vectashield mounting media (Clinisciences). Microscopy was performed using inverted microscope Leica DMI-6000. Images were taken with the HQ2 Coolsnap motorized by Metamorph software. All images were processed with ImageJ software.

#### Flow cytometry

Cells were trypsinized and resuspended in 1X Binding Buffer. 1*10^5 cells were incubated for 15 min at RT under lightless condition in 1X Annexin V binding buffer containing FITC Annexin V and 0.25 mg/μL propidium iodide (PI) according to FITC Annexin V Apoptosis Detection Kit (BD Biosciences). 10000 cells were sampled on BD FACSCaliburTM flow cytometer and cells were gated based on forward and side scatter. Fluorescence measurements were detected on the FL1 (Annexin V) and FL2 (PI). Data were analyzed using FlowJo software.

#### Immunoprecipitation (IP)

40 million cells were lysed in Buffer A (20 mM HEPES pH 7, 0.15 mM EDTA, 0.15 mM EGTA, 10 mM KCl), 10% NP40 and SR buffer (50 mM HEPES pH 7, 0.25 mM EDTA, 10 mM KCl, 70% (m/v) sucrose) supplemented with protease inhibitor (Sigma), 1 mM DTT and spermidine-spermine (0.15 mM each) to limit nuclei leak. Cell lysates were centrifuged at 2000 g for 5 min. The nuclei pellets were resuspended in sucrose buffer (20 mM Tris pH 7.65; 60 mM NaCl; 15 mM KCl; 0.34 M Sucrose) and high salt buffer was added (20 mM Tris-HCl pH 7.65; 0.2 mM EDTA; 25% glycerol; 900 mM NaCl; 1.5 mM MgCl_2_) to a final NaCl concentration of 300 mM, and incubated for 30 min in ice. Then, sucrose buffer was added to a final NaCl concentration of 100 mM. The nuclear extracts were treated with Micrococcal nuclease (0.0125 U/ml) and 1 mM CaCl_2_ at 37 °C during 15 min. Then, EDTA was added to a final concentration of 4 mM, and subsequent sonication for 5 min (15 sec ON, 1 min OFF) at medium frequency (Bioruptor Diagenode) was performed. The lysates were ultracentrifuged at 40000 rpm for 30 min and pre-cleared with protein G-agarose beads (Sigma) during 2 h, at 4°C. For immunoprecipitation using Myc-Trap technology (ChromoTek), the Myc-Trap beads were equilibrated with Wash buffer (10 mM Tris pH 7.5, 150 mM NaCl, 0.15 mM EDTA). Myc immunoprecipitations were carried out 1 h, at 4°C using 25 μL of Myc-Trap bead slurry and 500 μg of nuclear extract. The immunocomplexes were washed four times in Wash buffer and the proteins were eluted in NuPAGE® LDS Sample Buffer (Life Technologies) and 10X reducing agent at 96°C during 10 min. Finally, the immunoprecipitants were tested by western blot.

#### Mass spectrometry

##### Material

MS grade Acetonitrile (ACN), MS grade H2O, MS grade formic acid (FA), TRIS buffer, Tris(2-Carboxyethyl) Phosphine Hydrochloride (TCEP-HCl) and S-Methyl methanethiosulphonate (MMTS) were from ThermoFisher Scientific (Waltham, MA, USA). Chymotrypsin was from Promega (Madison, WI, USA). Ammonium bicarbonate (NH_4_HCO_3_) was from Sigma-Aldrich (Saint-Louis, MO, USA).

##### Samples preparation prior to LC-MS/MS analysis

5-minutes-cycles alternating NH_4_HCO_3_ 50mM and ACN were performed to destain gel plugs.

Proteins were reduced with 10 mM TCEP-HCl and alkylated with 20 mM MMTS. Then the sample was in gel-digested overnight with chymotrypsin at 37°C in a 100 mM TRIS buffer (50 µL final volume and add of 0.5 µg chymotrypsin). The digested peptides were loaded and desalted on Evotips provided by Evosep (Odense, Denmark) according to manufacturer’s procedure before LC-MS/MS analysis.

##### LC-MS/MS acquisition

Samples were analyzed on a timsTOF Pro 2 mass spectrometer (Bruker Daltonics, Bremen, Germany) coupled to an Evosep one system (Evosep, Odense, Denmark) operating with the 30SPD method developed by the manufacturer. Briefly, the method is based on a 44-min gradient and a total cycle time of 48 min with a C18 analytical column (0.15 x 150 mm, 1.9 µm beads, ref EV-1106) equilibrated at 40°C and operated at a flow rate of 500 nL/min. H_2_O/0.1 % FA was used as solvent A and ACN/ 0.1 % FA as solvent B.

The timsTOF Pro 2 was operated in PASEF mode1 over a 1.3 sec cycle time. Mass spectra for MS and MS/MS scans were recorded between 100 and 1700 m/z. Ion mobility was set to 0.75 - 1.25 V·s/cm2 over a ramp time of 180 ms. Data-dependent acquisition was performed using 6 PASEF MS/MS scans per cycle with a near 100% duty cycle. Low m/z and singly charged ions were excluded from PASEF precursor selection by applying a filter in the m/z and ion mobility space (Meier et al., 2015). The dynamic exclusion was activated and set to 0.8 min, a target value of 16000 was specified with an intensity threshold of 1000. Collisional energy was ramped stepwise as a function of ion mobility.

##### Data analysis

MS raw files were processed using PEAKS Studio (build 11.5, Bioinformatics Solutions Inc.). Data were searched against the Mus Musculus Swiss-Prot database (downloaded 2024_01, 17,144 entries). Parent mass tolerance was set to 20 ppm, with fragment mass tolerance at 0.05 Da. Semi-specific chymotrypsin cleavage was selected and a maximum of 2 missed cleavages was authorized. For identification, the following post-translational modifications were included: oxidation (M), deamidation (NQ), acetylation (K, Protein N-term), methylation (KR), dimethylation (KR), trimethylation (KR) as variables and beta-methylthiolation (C) as fixed. Identifications were filtered based on a 1% FDR (False Discovery Rate) threshold at both peptide and protein group levels. A minimum A-score of 10 was used to filter modified peptides based on the confidence in the position of the modified site. The A-score calculates an ambiguity score as -10 × log10(p), where the *p-*value indicates the likelihood that the peptide matches by chance.

#### Western Blot (WB)

Nuclear extracts or immunoprecipitants were resolved on pre-cast polyacrylamide gel cassettes (NuPAGE® Novex® 4-12% Bis-Tris) (Invitrogen) and 1X NuPAGE MES SDS Running Buffer and transferred into nitrocellulose membrane (Amersham) in 20 mM phosphate transfer buffer (pH 6.7). Membrane was blocked in 5% skim milk in PBST Buffer (1X PBS, 0.2% Tween 20) and incubated overnight at 4°C with the primary antibodies against Myc (Cat# sc40; Santa Cruz Biotechnologies), pan-tri-methyl-lysine (Cat# 14680; Cell Signaling Technology), Histone 3 (Cat# sc-8654; Santa Cruz Biotechnologies), HP1γ (Cat# 05-690; Millipore). Membranes were incubated with the appropriate secondary antibody coupled to HRP, revealed using West Dura kit (Pierce, Rockford, USA) and ChemiSmart 5000 system (Vilber Lourmat). Quantification of western blot signals was performed using Image J software.

To quantify protein levels in Western blot images, a ratio-based analysis of the immunoprecipitation data was performed. This involved measuring the intensity of each protein band and normalizing it to the corresponding loading control band.

#### Chromatin immunoprecipitation (ChIP)

Cells were cross-linked directly in the culture plate with 1% formaldehyde (culture medium supplemented with 1% formaldehyde (Sigma), 15 mM NaCl, 0.15 mM EDTA, 0.075 mM EGTA, 0.015 mM HEPES pH 8) during 10 min at RT. Formaldehyde was quenched with 0.125 M glycine and cells were washed in PBS and pelleted. Cells were then incubated on the wheel at 4°C for 10 min in Buffer 1 (50 mM HEPES /KOH pH 7.5; 140 mM NaCl; 1 mM EDTA; 10% Glycerol; 0.5% NP-40; 0.25% Triton X-100). After centrifugation, cells were incubated on the wheel at 4°C for 10 min in Buffer 2 (200 mM NaCl; 1 mM EDTA; 0.5 mM EGTA; 10 mM Tris pH 8.0). Nuclei were then pelleted by centrifugation, resuspended in Buffer 3 (50 mM Tris pH 8.0; 0.1% SDS; 1% NP-40; 0.1% Na-Deoxycholate; 10 mM EDTA; 150 mM NaCl), and sonicated for 20 min (30 sec ON, 30 sec OFF) (Bioruptor Diagenode), yielding genomic DNA fragments with a bulk size of 150-600 bp. All buffers were supplemented with protease inhibitors prior to usage. Chromatin corresponding to 5 μg (for ChIP H3K9me3) or 10μg (for ChIP Flag and HP1γ) of DNA was pre-cleared with protein G-agarose beads (Sigma) during 2h at 4°C. 1% of chromatin extracts were taken aside for inputs. Immunoprecipitations with 3 μg of anti-H3K9me3 (Cat# ab8898; Abcam), 5 μg of anti-Flag (Cat# F7425; Sigma-Adrich) or 5 μg of anti-HP1γ (Cat# 05-690; Millipore) antibodies were carried out overnight at 4°C. Pierce™ Protein A/G UltraLink™ Resin (Thermo-Fisher Scientific) was blocked overnight at 4°C with 0.3% BSA. Immune complexes were recovered by adding pre-blocked protein A/G UltraLink™ beads and incubated for 2 h, at 4°C. Beads were washed twice with Low salt buffer (0.1% SDS; 1% Triton; 2 mM EDTA; 20 mM Tris pH 8.0; 150 mM NaCl), twice with High salt buffer (0.1% SDS; 1% Triton; 2 mM EDTA; 20 mM Tris pH 8.0; 500 mM NaCl), once with LiCl wash buffer (10 mM Tris pH 8.0; 1% Na-deoxycholate; 1% NP-40, 250 mM LiCl; 1 mM EDTA), and once with TE supplemented with 50 mM NaCl. Cross-linked chromatin was eluted in TE with addition of 1% SDS and 0.2 M NaCl at 65°C during 45 min. ChIP-enriched samples and inputs were then reverse cross-linked at 65°C overnight and treated with 0.3 μg/mL RNase A. Eluted material was incubated with Proteinase K 2 h at 37°C. DNA was then obtained using the kit MinElute PCR Purification Kit (Qiagen) and DNA was resuspended in water. In order to improve the enrichment for ChIP-Flag and ChIP-HP1γ, five independent immunoprecipitations were performed simultaneously and the DNA was concentrated using the kit DNA Clean & Concentration^TM^ (Zymo Research). qPCR was performed using PowerSYBR Green PCR Master mix (Applied Biosystems) and analyzed on a Via 7 System (Applied Biosystems). ChIP-qPCR results were represented as a percentage (%) of IP/input signal (% input).

#### ChIP-sequencing (ChIP-seq)

DNA and input replicate samples from two independent time course experiments were sequenced at Platform GENOM’IC - Institute Cochin. The immunoprecipitated DNA was quantified using the Qubit™ dsDNA High Sensitivity HS assay (ThermoFisher Scientific) and fragment size distribution (between 100-600 bp) was analyzed by capillary electrophoresis (Agilent 2200 TapeStation Nucleic Acid System with the High sensitivity D5000 kit). Libraries were prepared using the MicroPlex V2 library preparation™ kit (Diagenode) following the manufacturer’s instructions. Starting amount of fragmented DNA varied between sub-nanograms and 500 pg of DNA (from ChIP Flag-SETDB1), 1.5 ng of DNA (from ChIP H3K9me3) and 1.5 ng for inputs were used. After end repair of the double-stranded DNA templates, we ligated the cleavable stem-loop adaptors and amplified the adaptor-enriched DNA with high fidelity amplification of 10 PCR cycles to add Illumina compatible indexes. The libraries were then purified with the Agencourt AMPure XP bead-based purification system. Final libraries were quantified with Qubit™ dsDNA HS assay (ThermoFisher Scientific) and size distributions were monitored by capillary electrophoresis with Agilent 2100 Bioanalyzer using High Sensitivity™HS kit (Agilent). After pooling, libraries were paired-end sequenced (2X40 bp) on a Nextseq 500 instrument (Illumina).

#### ChIP-seq data analysis

Paired-end ChIP-seq reads (2 × 40 bp) were aligned to the Mus musculus reference genome (mm10) using Bowtie2 (v2.5.3) with default parameters. Reads overlapping with blacklisted regions (ENCFF547MET) were excluded to reduce potential artifacts.

Signal tracks were generated using DeepTools2 bamCompare (v3.5.4), calculating the ratio of RPKM-normalized immunoprecipitated (IP) signal to input across 50 bp bins with a pseudocount of “0 1”. The resulting BigWig files were visualized using the Integrative Genomics Viewer (IGV). Peak calling for FLAG-SETDB1 ChIP-seq replicates was performed using MACS2 (v2.2.9.1) with q-value<0.05 and broad option. Overlapping peaks between biological replicates were filtered and merged using BEDTools (v2.31.1) intersect and mergeBED. Merged peaks were annotated using the ChIPSeeker R package (1.42.1) within the RStudio environment (2024.12.1 Build 563, R version 4.4.3). Repeat annotations were assigned using RepeatMasker data for the mm10 genome.

To visualize ChIP-seq signal at specific genomic loci, replicate alignments were consolidated using Picard MergeBamFiles (v1.56.0) to produce a unified BAM file for downstream analysis. For SETDB1 and H3K9me3 datasets, DeepTools2 bamCompare (v3.5.4) was used to compute fold-change ratios (IP/Input, RPKM) and convert BAM files to BigWig format using 50 bp bins. Signal pile-ups were analyzed using DeepTools2 computeMatrix. For SETDB1 peaks, the reference-point mode was used to center signals. For IAP repeats, the scale-regions mode was applied with a “regionBodyLength” of 7000 bp. Heatmaps and signal profiles were generated using DeepTools2 plotHeatmap and plotProfile.

Repeat binding analysis was conducted using RepeatMasker annotations and BEDTools MultiCovBed (v2.31.1) on deduplicated, merged alignment files. RPKM values were calculated for both IP and input samples, and the IP/Input ratio was used to generate bar plots.

#### RNA reverse transcription and quantitative PCR (RT-qPCR)

Total RNA was extracted using RNeasy mini-kit (Qiagen) following manufacturer’s procedures. DNase (Qiagen) treatment was performed to remove residual DNA. With High-Capacity cDNA Reverse Transcription Kit (Applied Biosystems), 1 μg of total RNA was reverse transcribed. Real-time quantitative PCR was performed to analyze relative gene expression levels using SYBR Green Master mix (Applied Biosystems) following manufacturer indications. Relative expression values were normalized to the housekeeping genes mRNA *Cyclophilin A* or *GAPDH*.

#### RNA-sequencing (RNA-seq)

RNA was isolated as described above. Three independent biological replicates were sequenced per cell condition at Platform GENOM’IC - Institute Cochin. After RNA extraction, RNA concentrations were obtained using a fluorometric Qubit RNA HS™ assay (ThermoFisher Scientific). The quality of the RNA (RNA integrity number 8.2) was determined on the Agilent 2100 Bioanalyzer (Agilent Technologies, Palo Alto, CA, USA) following the manufacturer’s instructions. To construct libraries, 1 μg of high-quality total RNA sample (RIN >8) was processed using Truseq® stranded total RNA kit (Illumina) according to the manufacturer’s instructions. Briefly, after removal of human ribosomal RNA (using Ribo-zero® rRNA) confirmed by QC control on pico chip™ on the Agilent 2100 Bioanalyzer (Agilent Technologies, Palo Alto, CA, USA), total RNA molecules were fragmented and reverse-transcribed using random primers. Replacement of dTTP by dUTP during the second-strand synthesis permitted the achievement of the strand specificity. Addition of a single A base to the cDNA was followed by the ligation of adapters. Libraries were quantified by qPCR using the KAPA Library Quantification Kit for Illumina Libraries (KapaBiosystems) and library profiles were assessed using the DNA High Sensitivity™HS kit on an Agilent Bioanalyzer 2100. Libraries were sequenced on an Illumina® Nextseq 500 instrument using 2x40 base-lengths read V2 chemistry in a paired-end mode.

#### RNA-seq data analysis

RNA-seq analysis was conducted using RASflow_EDC v1.3 (https://github.com/parisepigenetics/RASflow_EDC), an in-house pipeline adapted from the original RASflow framework (Zhang and Jonassen, 2020). Briefly, raw sequencing reads underwent quality control and trimming to remove low-quality bases and adapter sequences.

Reads were aligned to the Mus musculus reference genome (*mm10*) using STAR (Dobin et al., 2013). Gene-level quantification was performed with featureCounts (Liao et al., 2019), applying default parameters for gene annotation and the -M --fraction options for repeat elements. Annotation was based on GENCODE vM25.

Differential expression analysis was carried out using the DESeq2 R package (Love et al., 2014). Genes with an adjusted *p*-value < 0.05 were considered significantly differentially expressed.

All analyses were executed on the iPOP-UP HPC cluster, hosted by RPBS and supported by Université Paris Cité (IDEX).

#### Downstream Analysis of Differentially Expressed Genes and Repeats

The complete code for downstream analysis of differentially expressed genes (DEGs) and repeat elements is publicly available and maintained under full version control in the GitHub repository of the Paris Epigenetics Unit: https://github.com/parisepigenetics/RASflow_EDC.

Differentially expressed genes were filtered based on log₂ fold-change thresholds and selected for downstream analysis using R packages. Hierarchical clustering was performed using the complete linkage method to generate heatmaps via the ComplexHeatmap R package.

Gene Ontology (GO) enrichment analysis was conducted using the clusterProfiler R package, and results were visualized as dot plots to highlight significantly enriched biological processes.

The overlap between upregulated genes in cells expressing either the automethylation-deficient (AA) or catalytic-dead (CA) SETDB1 mutants and genes associated with SETDB1- and H3K9me3-enriched genomic regions in wild-type (WT) SETDB1-expressing cells was visualized using an UpSet plot, generated with UpSet R package.

### STATISTICAL ANALYSES

Statistical analyses of the NGS data are included within the corresponding section of the STAR METHODS. In addition, the collected data was analyzed using Statistical Package for Social Sciences (SPSS Version 25, Chicago IL). Normality of the data was assessed using the Shapiro-Wilk test. Results were presented as mean and standard deviation (SD) or median and interquartile range (IQR). Differences between groups were compared using *t*-test or Mann Whitney’s unpaired test based on normality test. Statistical significance was assessed at the α <0.05 levels for RT-qPCR, ChIP-qPCR, cell survival apoptosis tests, and ratio-based analysis of immunoprecipitation.

---

**Figure S1.**
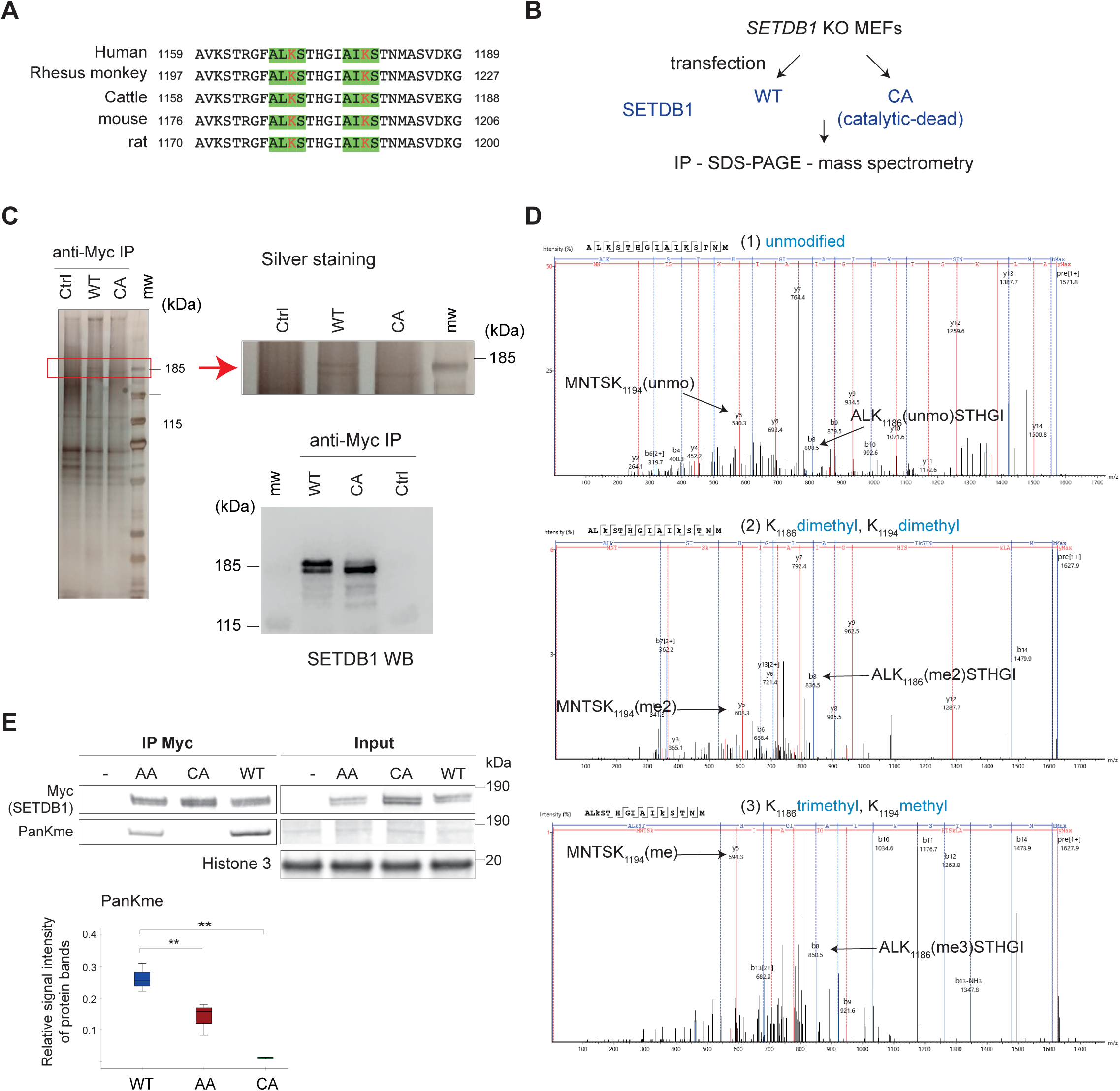
related to Figure 1. SETDB1 undergoes auto-methylation on lysines K1170 and K1178 in vitro and is methylated in the cells. **A.** Amino acid sequence alignment of a conserved region of SETDB1 protein across five species: Human, Rhesus monkey, Cattle, Mouse, and Rat. The alignment shows positions 1159–1189 (human numbering) with conserved amino acid residues highlighted. Green boxes indicate conserved A-R/L/I-KS motif. **B.** Diagram of the experimental setup used to study SEDTB1 methylation in iMEFs by mass spectrometry. **C.** Myc immunoprecipitation was performed on nuclear fractions from *Setdb1* KO iMEFs transfected with either an empty vector (Ctrl), wild-type SETDB1 (WT), or a catalytically inactive mutant (CA) expression vector. Immunoprecipitated proteins were visualized by silver staining (upper panel), and SETDB1 protein levels were confirmed by western blot analysis (lower panel). **D.** Representative mass spectra of the unmodified peptide, and of the two most abundant modified peptides, which included dimethylation at both K1186 and K1194 for one and trimethylation at K1186 and monomethylation at K1194 for the other one. Fragments b8 and y5 are common to all fragmentation spectra and allow the modifications to be positioned with certainty at K1186 and K1194, respectively. **E.** The catalytic activity of SETDB1 is required for its methylation in mouse cells. Three different Myc-tagged SETDB1 cDNAs, including full-length (WT), automethylation-deficient (AA), and catalytic-inactive (C1243A, named CA) forms, were cloned and transiently expressed in *Setdb1* KO MEFs. Myc-SETDB1 was immunoprecipitated by Myc-Trap technology (IP Myc) and subjected to Western blot analyses using anti-Pan-tri-methyl-lysine (PanKme) and anti-Myc antibodies (left panel). Inputs, including histone H3 as a loading control, are shown (right panel). Protein levels were quantified and presented as the ratio between the PanKme signal over the Myc-tagged SETDB1 (lower panel). Statistical significance was determined using the unpaired Mann–Whitney test (n = 3 biological replicates). **: p < 0.05.

**Figure S2.**
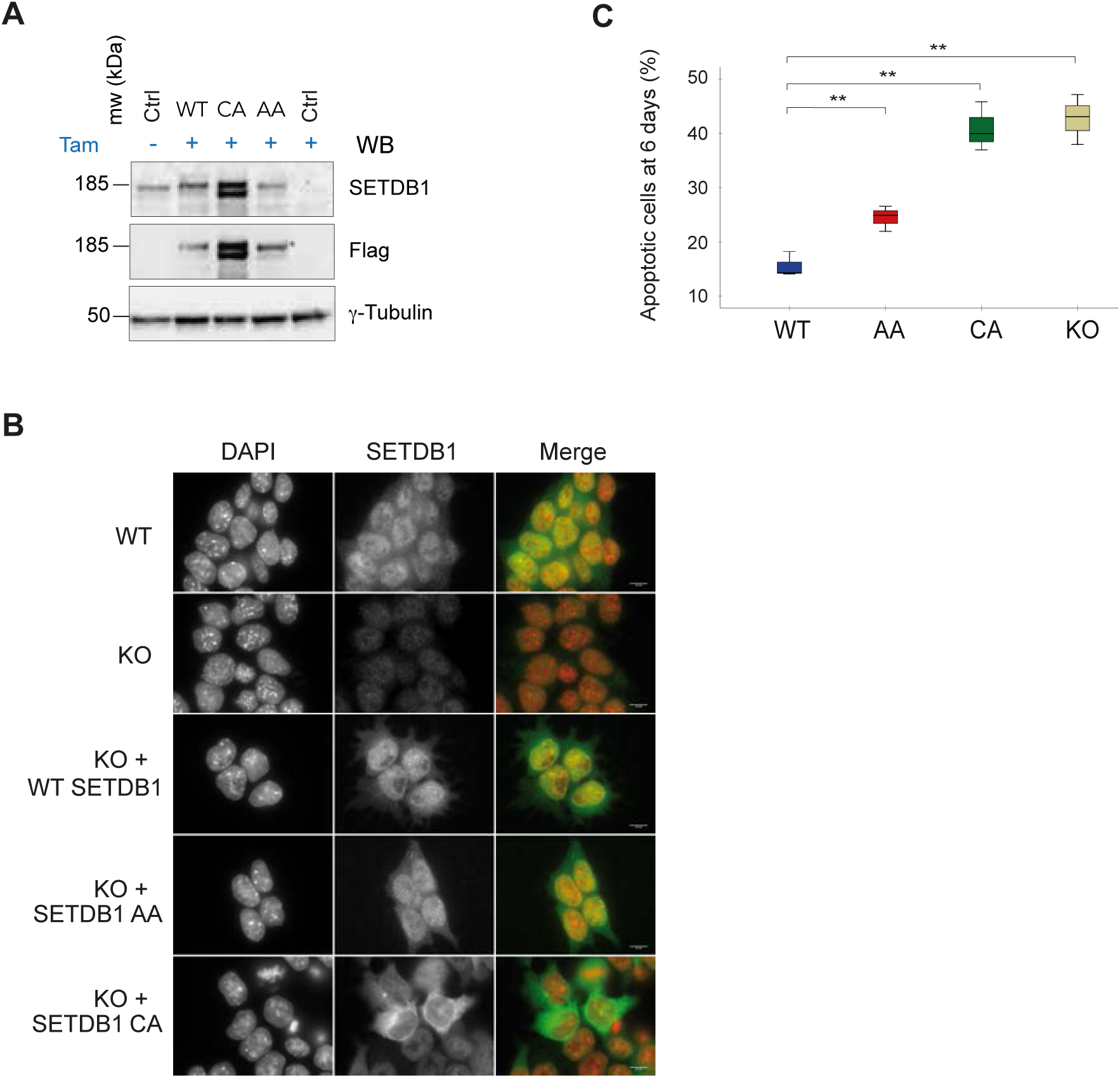
related to Figure 2. The automethylation-deficient SETDB1 mutant is enzymatically active, but it compromises mESCs’ fitness. **A-B.** Stable expression of Flag-tagged SETDB1 forms in TT2 mESCs harboring floxed *Setdb1* and stably transfected with the empty vector (Ctrl) or expression vectors for wild-type (WT), automethylation-deficient (AA) or catalytically inactive (CA) SETDB1. ESC lines were treated with Tamoxifen (TAM) for 96h to knock-out endogenous *Setdb1*. Exogenous and stably-expressed Flag-SETDB1 was revealed by Western blot (A) by using Flag and SETDB1 antibodies and γ−Tubulin as a loading control, or by indirect immunofluorescence (green) and DNA was labelled with DAPI (red) (B). Scale bar in (B) = 10 μm. **C.** The automethylation-deficient SETDB1 does not fully rescue the apoptosis increase observed after *Setdb1* KO in ESCs. ESCs lines described in (A) were treated with tamoxifen for 4 days, followed by 2 days without treatment, and apoptosis was determined by the percentage (%) of cells marked positive for Annexin V and/or Propidium iodide. Data followed normal distribution; consequently, Student *t*-test was applied to test statistical significance (n = 3 biological replicates) **: p<0.05.

**Figure S3.**
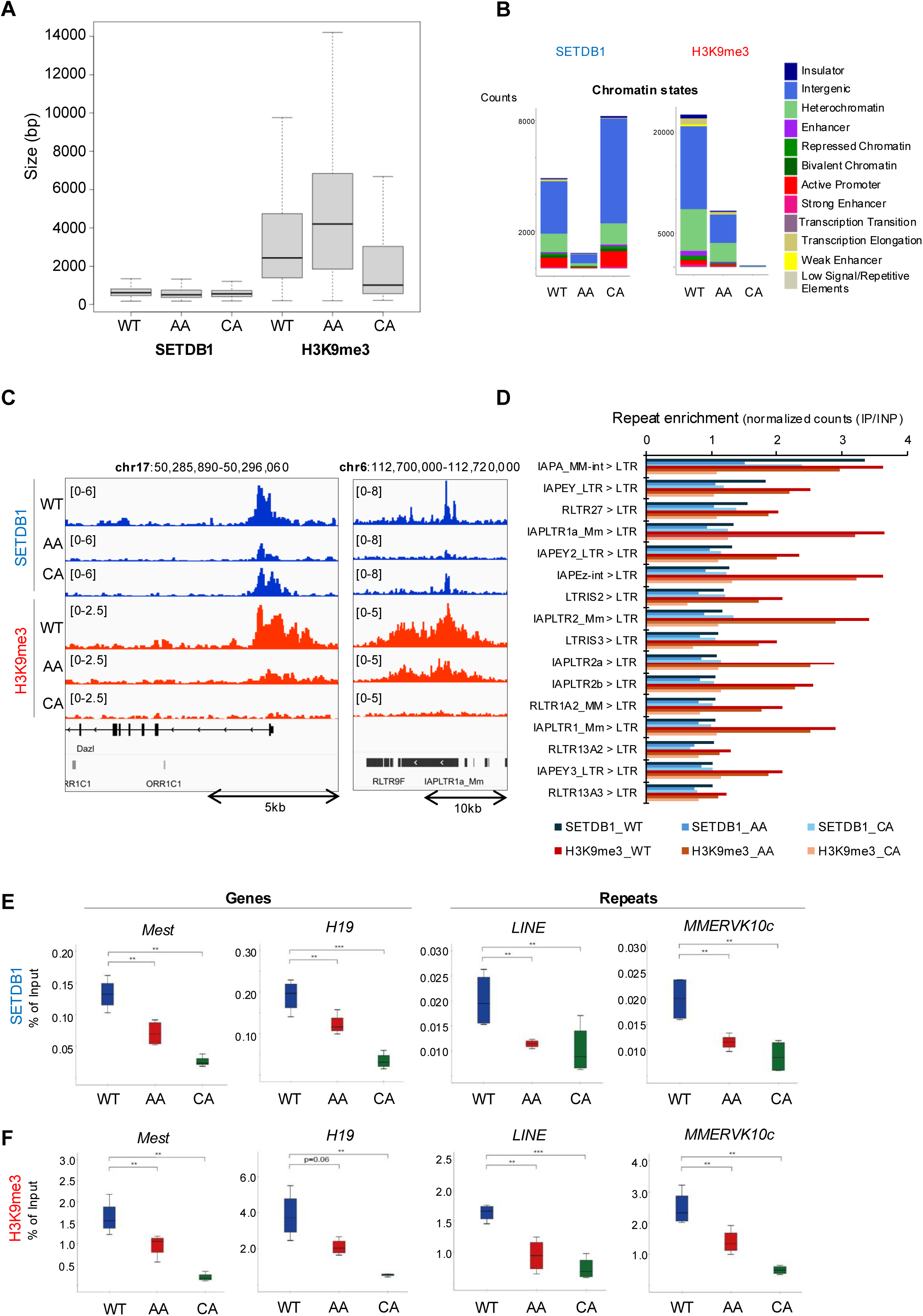
related to Figure 3. SETDB1 auto-methylation modulates its genomic binding capacity and influences the deposition of H3K9me3 at both coding genes and transposable elements. ChIP-seq and ChIP-qPCR data from *Setdb1* cKO mESCs re-expressing wild-type (WT) SETDB1, auto-methylation-deficient (AA) or catalytic-dead (CA) SETDB1 mutants. **A.** Boxplot showing the distribution of peak widths (in base pairs) identified by ChIP-seq for Flag-SETDB1 and H3K9me3 in mESCs expressing wild-type (WT), automethylation-deficient (AA), or catalytically inactive (CA) SETDB1 mutants. **B.** Genomic annotation of Flag-SETDB1 binding sites across chromatin state (https://github.com/guifengwei/ChromHMM_mESC_mm10, (Pintacuda et al., 2017)), based on ChIP-seq enrichment for Flag-SETDB1 and H3K9me3. Data shown represent enriched genomic regions consistently identified in both biological replicates. **C.** IGV genome browser tracks displaying ChIP-seq signal for Flag-SETDB1 and H3K9me3 at representative loci—including the *Dazl* gene, RLTR9, and IAPLTR1a_Mm repetitive elements—on chromosome 17 (chr17) and chromosome 6 (chr6) in mESCs expressing wild-type (WT), automethylation-deficient (AA), or catalytically inactive (CA) SETDB1 mutants. **D.** Bar plots showing fold changes (FC > 1) in ChIP-seq signal for individual long terminal repeated (LTR) elements, illustrating the loss of SETDB1 binding and H3K9me3 enrichment in automethylation-deficient (AA), or catalytically inactive (CA) SETDB1 across a representative set of endogenous retroviral (ERV) families. The Y-axis indicates fold enrichment of normalized ChIP read density relative to input. **E-F.** Crosslinked ChIP-qPCR analysis of Flag-tagged SETDB1 (E) and H3K9me3 (F) at selected genes and repeated elements in mESCs expressing wild-type (WT), automethylation-deficient (AA), or catalytically inactive (CA) SETDB1 forms. Statistical significance was assessed using Student’s *t*-test for normally distributed data, or Mann–Whitney’s unpaired test otherwise (n = 4 biological replicates). **p < 0.01; ***p < 0.001; NS: not significant

**Figure S4.**
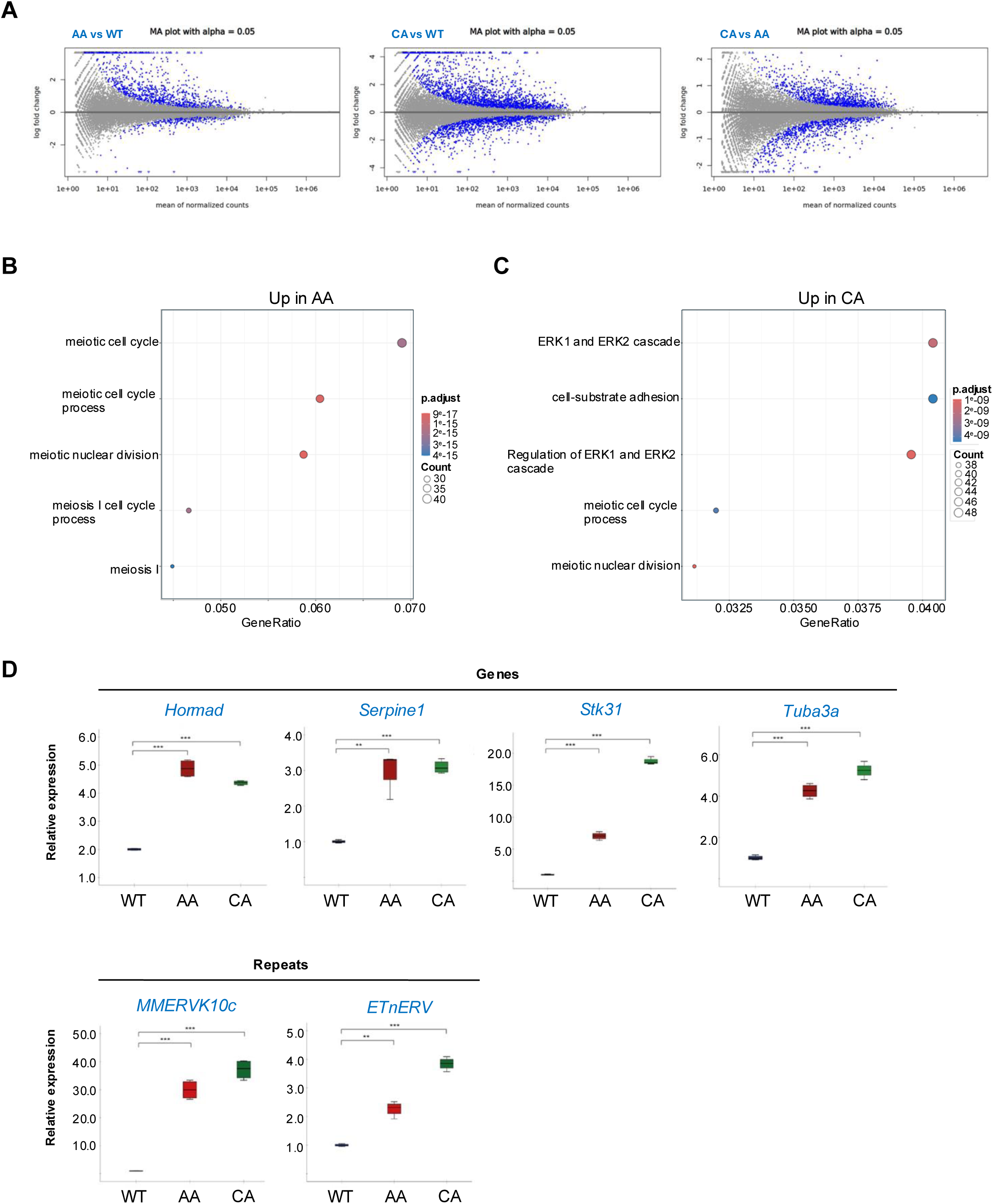
related to Figure 4. SETDB1 auto-methylation is required for its gene and repeat silencing activity. **A.** MA plot panels displaying differential gene expression across three comparisons: AA vs WT (left), CA vs WT (middle), and AA vs CA (right). Each plot shows log₂ fold change versus the mean of normalized counts. Significantly deregulated genes (p < 0.05) are highlighted in blue, with upregulated genes appearing above the horizontal line at zero and downregulated genes below. **B-C.** Dot plot showing enrichment of Gene Ontology (GO) biological processes for upregulated genes in mESCs expressing AA (B) or CA (C) SETDB1 mutants (fold change > 0.5; adjusted p-value < 0.05). The X-axis represents the GeneRatio, calculated as the number of upregulated genes divided by the total number of genes annotated to each GO term. Dot size reflects the number of genes associated with each process, while dot color indicates the adjusted p-value. The Y-axis lists the top enriched biological processes. **D.** RT-qPCR analysis of selected deregulated genes (upper panel) and repetitive elements (lower panel), as in Figure 4H, in *Setdb1* conditional knockout (cKO) mESCs re-expressing either wild-type (WT), automethylation-deficient (AA) or catalytically inactive SETDB1 mutants. mRNA levels were normalized to Cyclophilin A expression. Statistical significance was determined using Student’s t-test for normally distributed data or Mann–Whitney’s unpaired test otherwise (n = 4 biological replicates). Significance levels are indicated as **p < 0.01; ***p < 0.001

**Figure S5.**
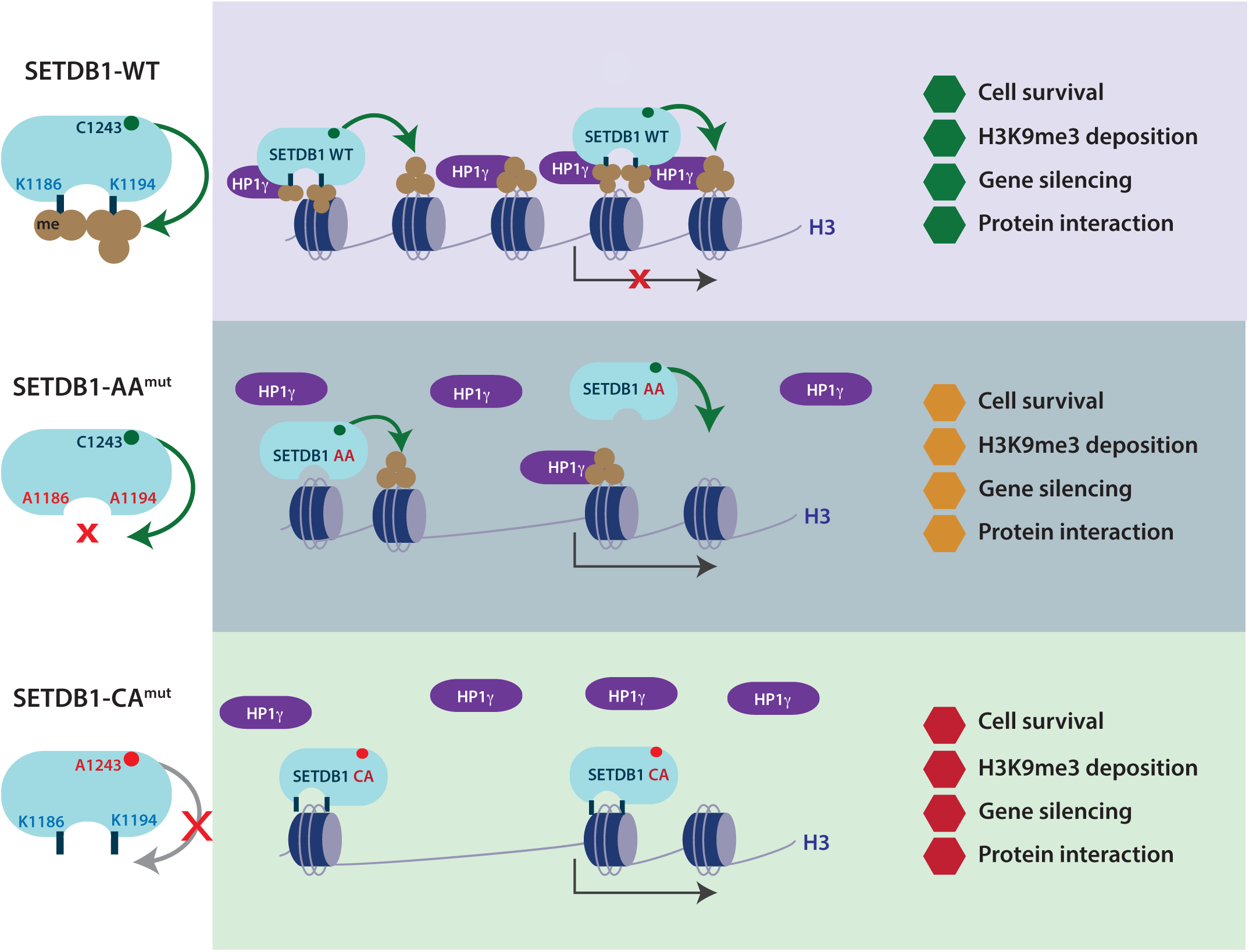
Working model. *Left:* Schematic representation of the SETDB1 variants used in this study, indicating their enzymatic activity and automethylation status at lysines K1186 and K1194. ***Right:*** Wild-type (WT) SETDB1 is recruited to its chromatin binding sites and stabilized through interactions with HP1γ and other chromodomain-containing proteins (not depicted), promoting efficient H3K9me3 deposition, gene and repeat silencing, and ESC survival (upper panel). The automethylation-deficient mutant (SETDB1-AA), although capable of depositing H3K9me3, binds less stably to chromatin due to weakened interactions with its chromodomain-containing partners. This results in reduced H3K9me3 levels, partial derepression of genes and repeats, and decreased cell viability (middle panel). The catalytically inactive mutant (SETDB1-CA) is recruited to chromatin but fails to interact with chromodomain-containing partners or deposit H3K9me3, thereby compromising gene and repeat repression and ESC survival (lower panel).

